# Antigen affinity and site of immunization dictate B cell recall responses

**DOI:** 10.1101/2024.04.19.590239

**Authors:** M. Termote, R. C. Marques, E. Hyllner, M. V. Guryleva, M. Henskens, A. Brutscher, I.J.L. Baken, X. Castro Dopico, A. Dalmau Gasull, B. Murrell, L. Stamatatos, L. S. Westerberg, P. Dosenovic

**Affiliations:** Department of Microbiology, Tumor and Cell Biology, Karolinska Institutet, 171 65 Solna, Sweden; Vaccine and Infectious Disease Division, Fred Hutchinson Cancer Research Center, Seattle, WA 98109, USA

**Keywords:** Memory B cells, Germinal center, B cell recall responses, sequential immunization, antigen affinity, germinal center refueling, secondary germinal centers, HIV-1 vaccine

## Abstract

Protective antibodies against HIV-1 require unusually high levels of somatic hypermutations (SHMs) introduced in germinal centers (GCs). To achieve this, a sequential vaccination approach was proposed. Using HIV-1 antibody knock-in mice with fate-mapping genes, we examined if antigen affinity affects the outcome of the B cell recall response. Compared to high affinity boost, low affinity boost resulted in decreased numbers of memory-derived B cells in secondary GCs, but with higher average SHM, indicating an affinity threshold for memory B cells to enter secondary GCs. Upon boosting local LNs, numbers of residual GC B cells increased independent on antigen affinity, while average SHM decreased. Our results demonstrate that antigen affinity and location of the boost affect the outcome of the B cell recall response. These results can help guide the design of vaccine immunogens aiming to selectively engage specific B cell clones for further SHM diversification.

## Introduction

Antibodies are the correlate of protection for most existing vaccines ^1^. However, to protect against variable pathogens, antibodies are required to express unusual qualities. For HIV-1, it is believed that a successful vaccine needs to elicit broadly neutralizing antibodies (bNAbs) that bind the more conserved epitopes of the HIV-1 envelope glycoprotein (Env) and exhibit high levels of somatic hypermutation (SHM) ^2–4^. These antibodies develop in a fraction of infected individuals ^5,6^ but are not readily elicited during vaccination, with the exception of a few rare studies in animal models ^7–9^. Data from prospective studies of HIV-1-infected individuals show that bNAb-B cell clones develop over time in response to the continuously mutating viruses ^10–13^. This finding, together with the fact that germline versions of bNAbs fail to bind native HIV-1 Env proteins ^14–18^, supported the idea of so called ‘germline-targeting’ sequential vaccination approaches for HIV-1, aimed to first target selected naïve B cell clones and then drive them through the process of GC SHM ^19^.

To test such strategies and to investigate the mechanism of eliciting bNAbs by sequential immunization in a systematic process *in vivo*, several laboratories generated human Ig knock-in mice, expressing the germline reverted VDJ/VJ-sequences of different HIV-1 bNAbs. Results from such immunization studies confirmed that HIV-1 Env-based germline-targeting antigens could activate naïve bNAb-precursor B cells, which could gain selected bNAb features by switching the initial immunogens to heterologous, more native-like Env antigens, in subsequent boosts ^20–27^. The HIV-1 germline-targeting antigen eOD-GT8, engineered to activate naïve B cells consisting of the VH1-2*02/03 germline gene of the VRC01-like family of bNAbs ^18,20^, has in addition shown potential in activating the desired precursor B cells in humans ^28^. The sequential vaccination approach against HIV-1 is based on the assumption that somatically hypermutated memory B cells that form in response to a prime immunization are recalled by sequential antigens to enter secondary GCs for further diversification upon boost ^29,30^. Alternatively, GC B cells activated in response to a prime immunization are re-stimulated, or ‘refueled’, by a local boost immunization to lengthen their time in the GC ^30^. To successfully design appropriate antigens that can effectively promote these events upon repetitive immunization, a deeper understanding of the B cell recall response is needed.

Studies in both mice and humans show that memory B cells formed upon immunization or infection are a heterogenous population of cells ^31,32^. Upon recall, memory B cells are stimulated to quickly differentiate into antibody-producing cells ^33–35^ or enter secondary GCs for further diversification of their BCRs ^36–39^. It was recently shown that most GC B cells forming upon boost originate from B cells with no previous GC experience and only a minority are derived from memory B cells ^34,40^. Furthermore, Mesin et. al demonstrated that recall GC and plasma cell responses are dominated by fewer than 10 memory B cell-derived clones. The regulation of the memory B cell recall response is incompletely understood but is thought to depend on intrinsic factors that correlate with the origin and phenotype of the memory B cell ^41^. In addition, it is likely affected by extrinsic factors such as the presence of pre-existing antibodies ^42–47^ and the presence of cognate T follicular helper (Tfh) cells ^41,47^, as well as the location of the memory B cells ^40^. However, whether antigen affinity plays a role in B cell recall responses, and if modifications of the antigen can dictate memory B cell fates, has not been evaluated.

To understand B cell recall responses to complex antigens in more detail, we performed prime/boost immunizations of a mouse model with both polyclonal-, and a knock-in (KI) BCR specific against a conserved epitope of HIV-1 Env. The KI B cells expressed fate-mapping genes which enabled us to visualize GC B cells and B cells emerging from GCs after prime immunization. Using Env antigens of different affinities to the KI BCR as immunogens, we show that the number of memory-derived B cells in secondary GCs can be increased using a high affinity antigen upon boost, but that a low affinity antigen selectively recruits more mature memory B cells with higher levels of SHM for GC entry. Further, we show that the refueling of GCs results in increased numbers of GC B cells originating from the primary response, with lower average levels of SHM. Our findings have implications for the design of sequential vaccine approaches against viruses such as HIV-1, where the elicitation of epitope-specific antibodies with high levels of SHM is critical.

## STAR Methods

### Mice and immunizations/treatments

The 3BNC60 SI (<u>s</u>ynthetic intermediate) mouse, generously provided by M. Nussenzweig, is a human antibody knock-in (KI) mouse expressing the pre-rearranged mature (somatically hypermutated) heavy chain (VDJ), and the predicted germline-reverted light chain (VJ) of the HIV-1 bNAb 3BNC60 ^17,48^. 3BNC60 SI KI B cells used in experiments were heterozygous for both Ig KI alleles. S1pr2-ERT2cre mice were generously provided by T. Kurosaki ^49^ and *Rosa26*-tdTomato mice (Ai14; Rosa-CAG-LSL-tdTomato-WPRE Stock No. 007914) were purchased from Jackson Laboratory. B6.SJL (CD45.1^+^) mice were obtained from Charles River Laboratories. Male and female mixed bone marrow chimera mice were generated by injecting 2-3 million bone marrow cells, of which ≈ 3% were IgH^3BNC60mut/+^, IgL^3BNC60gl/+^, S1PR2^ERT2cre/+^, RS26^lsl-tdTomato/+^ (CD45.2^+^) and ≈ 97% were polyclonal S1PR2^ERT2cre/+,^ RS26^lsl-tdTomato/+^ (CD45.1^+^/2^+^), into recipient B6.SJL mice which had previously been irradiated with a total dose of 10 Gy. 8 weeks after reconstitution, the ratio of the two genotypes was confirmed by analyzing blood lymphocytes for CD45. Unless indicated differently, mice were pre-primed with 40 μg 2W1S, an MHCII-restricted peptide which is immunogenic in C57BL/6 mice ^50,51^. At indicated time points, mice were immunized s.c. with 5 μg/footpad of soluble HIV-1 Env proteins (Env-N276D, wtEnv, Env-DMRS) in Addavax adjuvant (InvivoGen) according to manufacturer’s instructions. After prime immunization at indicated time points, activated B cells were fate-mapped by activation of the Cre recombinase in S1PR2-ERT2cre, by oral administration of 6-12 mg of tamoxifen (Sigma-Aldrich) in 200 μL corn oil (Sigma-Aldrich) at indicated time points. At indicated time points, serum was collected from the tail vein and mice were sacrificed for analysis. All animal experiments were performed according to the protocols described in ethical permits (11159/18 and 05294-2023) of L. Westerberg, approved by the Stockholm North Animal Ethics Committee (Stockholm, Sweden).

### Recombinant antigen and monoclonal antibody expression and purification

The 3BNC60 antibody belongs to the VRC01-family of HIV-1 bNAbs and is composed of the human germline genes VH1-2*02 and VK1D-33 and binds the CD4 binding site (CD4bs) of Env gp140 with high affinity (6.81 x 10^-9^ M for YU2 Env) ^17^. The mutated VJ of 3BNC60 has evolved to accommodate the N-linked glycosylation at position N276 of HIV-1 Env. Hence, the VJ germline reversion of 3BNC60 SI results in a lower affinity binding to Env (1.93 x 10^-4^ M for 426c Env) ^48^. To study the role of affinity for B cell recall responses using the 3BNC60 SI KI mouse model, in addition to 426c Env (wtEnv) we also used HIV-1 Env proteins which bind to 3BNC60 SI with higher affinities, including an Env protein where the glycan at N276 has been removed (Env-N276D; 3.87 x 10^-5^ M for 426c Env) and an Env protein with a different combination of mutations that eliminate the N276 NLGS and other NLGS in the V5 (S278R, N460D, N463D, G471S) referred to here as Env-DMRS (TM4, elsewhere; 1.06 x 10^-8^ M for 426c Env) ^15,48,52,53^. All Env constructs used were based on HIV-1 426c clade C, and unless indicated differently, conjugated to 2W1S. Env proteins were expressed from pTT3-vectors with or without an AviTag for subsequent biotinylation. pTT3-426c-gp140, pTT3-426c-N276D-gp140, pTT3-426c-DMRS-gp140, pTT3-426c-gp140-2W1S, pTT3-426c-N276D-gp140-2W1S, pTT3-426c-DMRS-gp140-2W1S and pcDNA3.1-eOD-GT8-his were transfected into 293F cells at a density of 10^6^ cells ml^−1^ in Freestyle 293 media (Gibco) using PEI MAX (Polysciences). Expression was performed in Freestyle 293 media for 6-7 d with shaking at 37 °C in 5% CO2. Cells and cellular debris were then removed by centrifugation at 4000 x *g* followed by filtration of the supernatant through a 0.2 µm filter. For Env proteins expressed from the pTT3-vectors, the supernatant was passed over an Agarose-bound Galanthus Nivalis Lectin (GNL) resin (Vector Laboratories), pre-equilibrated with GNL binding buffer (20 mM Tris, 100 mM NaCl, 1 mM EDTA; pH 7.4), followed by washing with GNL binding buffer. Bound protein was eluted with GNL binding buffer containing 1 M methyl α-D-mannopyranoside. The supernatant containing eOD-GT8-his, was passed over a nickel nitrilotriacetic acid (Ni-NTA) resin (Thermo Scientific) pre-equilibrated with wash buffer (20 mM imidazole in PBS, pH 7.4), followed by washing with wash buffer, and then eluted with elution buffer (250 mM imidazole in PBS, pH 7.4). All proteins were further purified using a 16/600 size-exclusion column (SEC) (Cytiva) pre-equilibrated in PBS. Fractions containing trimeric gp140 protein or monomeric eOD-GT8 were pooled and concentrated. AviTagged Env variants were biotinylated *in vitro* using BirA500 (Avidity) according to the manufacturer’s instructions, followed by dialysis of the proteins in PBS using 100K MWCO Dialysis Cassettes (Thermo Scientific) to remove residual unligated biotin and BirA enzyme. Monoclonal antibodies were produced by co-transfecting the appropriate heavy and light chain expression plasmids at a 1:1 ratio into 293F cells as described above. Clarified supernatants were applied to Protein G Sepharose 4 Fast Flow (Cytiva) followed by washing with PBS. Antibodies were eluted in 0.5 ml fractions using 0.1 M Glycine pH 2.2 into 1.5 ml centrifuge tubes containing 0.2 ml of 1 M Tris-HCl pH 9.1. Fractions containing protein were pooled and buffer-exchanged into PBS using 20K MWCO Dialysis Cassettes (Thermo Scientific). Proteins and antibodies were aliquoted and frozen at −80°C.

### ELISA

High-binding 96-well plates (Corning) were coated with 0.1 µg of indicated protein per well. After incubation overnight at 4 °C, plates were washed 3 times in wash buffer (PBS with 0.05% TWEEN 20 (Sigma-Aldrich)) and incubated in blocking buffer (PBS with 2% milk) for 2 h. Serum samples and mAbs were added at indicated dilutions or concentrations and incubated for 2 h at room temperature. Plates were washed in wash buffer and HRP conjugated secondary antibody; anti-mouse IgG or anti-human IgG (Jackson Immuno Research), was added and incubated for 1 h 30 min at room temperature. Plates were washed and ABTS solution was added (Invitrogen). Absorbance was measured at 405 nm using a BioTek Eon Microplate Spectrophotometer.

### Flow cytometry and single B cell sorting

For flow cytometry and single B cell sorting, single cell suspensions were obtained by forcing lymph nodes through 70 μm filters into FACS buffer (2% FCS and 1 mM EDTA in PBS). Fc-receptors were blocked using rat anti-mouse CD16/32 (clone 2.4G2; BD). Cells were stained with fluorophore-conjugated antibodies including: anti-mouse/anti-human B220, anti-mouse/anti-human GL7, anti-mouse IgG1, anti-mouse CD138, anti-mouse CD80 (BioLegend), anti-mouse/anti-human B220, anti-mouse CD45.1, anti-mouse CD45.2, anti-mouse CD38, anti-mouse IgM (Invitrogen), anti-mouse CD45.1, anti-mouse CD45.2, anti-mouse CD38, anti-mouse IgM, anti-mouse IgG2a/b and anti-mouse PDL2 (BD). Env-binding B cells were stained using biotinylated AviTagged monomeric 426c Env-DMRS, Env-N276D and wtEnv (5 µg/ml) and detected using Streptavidin-PE (BD). Dump staining using antibodies specific for CD4, CD8, NK1.1, Gr1 and F4/80 (Invitrogen) were included in all flow cytometry stainings. Dead cells were excluded from analyses using Zombie NIR Fixable Viability Kit (BioLegend). Total number of cells were calculated using AccuCheck Counting Beads according to the manufacturer’s description (Invitrogen). Samples were acquired on a Fortessa (BD) and analyzed using FlowJo software (BD). tdT^+^ and tdT^-^ CD45.2^+^ B cells with either a GC B cell phenotype (CD4^−^, CD8^−^, Gr-1^−^, F4/80^−^, NK1.1^−^, B220^+^, GL7^+^, CD38^-^) or a memory B cell phenotype (CD4^−^, CD8^−^, Gr-1^−^, F4/80^−^, NK1.1^−^, B220^+^, GL7^-^, CD38^+^) were single cell sorted into 96-well plates using a FACSAria Fusion sorter (BD). Single sorted cells were lysed in 4 µl 0.5 X PBS (Gibco), 12 U RiboLock (Thermo Scientific) and 10 mM DTT (Invitrogen). Plates were stored at −80 °C until further processing. BCRs of single sorted cells were sequenced and amplified as previously described ^54,55^. The knock-in VJ sequences (3BNC60 LC) were sequenced and amplified from cDNA using the following primer pairs in nested PCR; 1_3BNC60_F_HK: 5’-GGGATGGTCATGTATCATCCTTTTTCTAG-3’ with 1mRK: 5’- ACTGAGGCACCTCCAGATGTT-3’ and 2_3BNC60_F_HK: 5’- GTAGCAACTGCAACCGGTGTACATTCT-3’ with 2mRK: 5’-TGGGAAGATGGAT ACAGTT-3’ ^53^. The knock-in VDJ sequences (3BNC60 HC) were sequenced and amplified from cDNA using the following primer pairs in nested PCR; 1_3BNC60_F_HK 5’- GGGATGGTCATG TATCATCCTTTTTCTAG-3’ with 1mRG 5’- AGAAGGTGTGCACACCGCTGGAC-3’ or Cµ outer 5’- AGGGGGCTCTCGCAGGAGACGAGG-3’ and 2_3BNC60_F_HK: 5’- GTAGCAACTGCAACCGGTGTACATTCT-3’ with 2mRG: 5’- GCTCAGGGAARTAGCCCTTGAC-3’ or Cµ inner 5’- AGGGGGAAGACATTTGGGAAGGAC-3’ ^25^. 2mRK and 2mRG or Cµ inner were used to sequence the VJ- and the VDJ products, respectively, by sanger sequencing.

### Sequence analysis

Light chain BCR sequences were truncated after amino acid position 90 (using sequential coordinates; ie. at the caron in DIATYYCQQ̌YEF in the germline reference sequence) to exclude regions with poor Sanger quality indicators. Phylogenies were inferred using FastTree2 (version 2.1.11, compiled with the double-precision flag to allow short branch lengths), under the generalized time-reversible nucleotide model, and visualized using FigTree.

### Statistical analysis

All data were analyzed using Prism 9 (GraphPad). Normal distribution was determined using the D’Agostino-Pearson normality test. Data that displayed normal Gaussian distribution were further analyzed for significant differences using two-tailed unpaired or paired Student’s t tests. Data that did not display normal Gaussian distribution were analyzed for significant differences using the Mann-Whitney U test or the Wilcoxon matched-pairs test. Data were considered significant at *, P ≤ 0.05; **, P ≤ 0.01; ***, P ≤ 0.001; ****, P ≤ 0.0001.

## Author contributions

M. Termote and P. Dosenovic planned and performed experiments, analyzed data, and wrote the manuscript. R. C. Marques, E. Hyllner, M. Henskens, A. Brutscher and I.J.L. Baken performed experiments, analyzed data, and provided scientific input. X. Dopico and A. Dalmau Gasull performed experiments. M. V. Guryleva and B. Murrell performed bioinformatic analysis of sequence data and provided scientific input. L. Stamatatos provided scientific input on experiments. All experimental work with mice was performed under the ethical permits of L. Westerberg, who also provided scientific input on experiments and provided technical assistance.

## Results

To enable the analysis of how antigen affinity affect B cell recall responses, we used mixed bone marrow chimera (MBMC) mice comprising an average of 1.8% antibody knock-in (KI) B cells expressing a fixed BCR, the 3BNC60 synthetic intermediate (SI) antibody (Supplementary Figure 1A) ^48^. B cells with this genetic background are here referred to as ‘KI B cells’ and bind the CD4 binding site (CD4bs) of HIV-1 Env proteins harboring different mutations, with increasing affinities. This was reflected by the binding of heterozygous KI B cells (IgH^3BNC60mut/+^, IgL^3BNC60gl/+^) to wtEnv, Env-N276D and Env-DMRS with increasing mean fluorescence intensity (MFI) (Supplementary Figure 1B) ^48^. Env-N276D and Env-DMRS are 99.9% and 99.40% identical to wtEnv in the amino acid sequence and even though they are modified to bind KI B cells with different affinities they induced an overall similar polyclonal GC B cell and antibody serum response at 15 days after immunization of wild type (WT) C57BL/6 mice (Supplementary Figure 2 A and B).

To facilitate detection of KI memory B cells after immunization, KI B cells expressed S1pr2-ERT2cre-tdTomato ^49^, which allows for tamoxifen-inducible tdTomato-labelling of germinal center (S1PR2+) KI B cells via Cre-recombination of the *Rosa26* allele. Due to the irreversible nature of the labelling, GC B cells that are successfully labelled will stay fate-mapped with tdTomato as tamoxifen-treatment is interrupted and the cells exit the GCs as antibody-secreting cells and memory B cells. Early, non-GC derived memory B cells will likely stay unlabeled after tamoxifen treatment due to the low or no expression of S1PR2 in the pre-GC stage ^56^.

### Immunization with intermediate affinity antigen and subsequent tamoxifen-treatment induces KI GC B cells of high labelling efficiency

To first determine the extent of activation and fate-mapping of KI GC B cells after immunization, MBMC mice were immunized with intermediate affinity Env-N276D in the two left foot pads. Mice were treated with tamoxifen on day 3, 6, 10, 17, 24, 31 and 38 post immunization. The time points were chosen to increase the chance of labeling cells that were activated both early-, during peak-, and later in the immune response. GC B cell formation was analyzed in draining- and non-draining LNs 10, 21, 35 and 42 days after immunization (Figure 1A). When comparing total GC B cells in Env-N276D-immunized and control non-immunized (adjuvant only) mice, we detected modest differences with a statistical increase in the draining LNs only on day 10 after Env-N276D-immunization (Figure 1B, Supplementary Figure 3A: gating strategy 1 and Supplementary Figure 3B).

**Figure 1:**
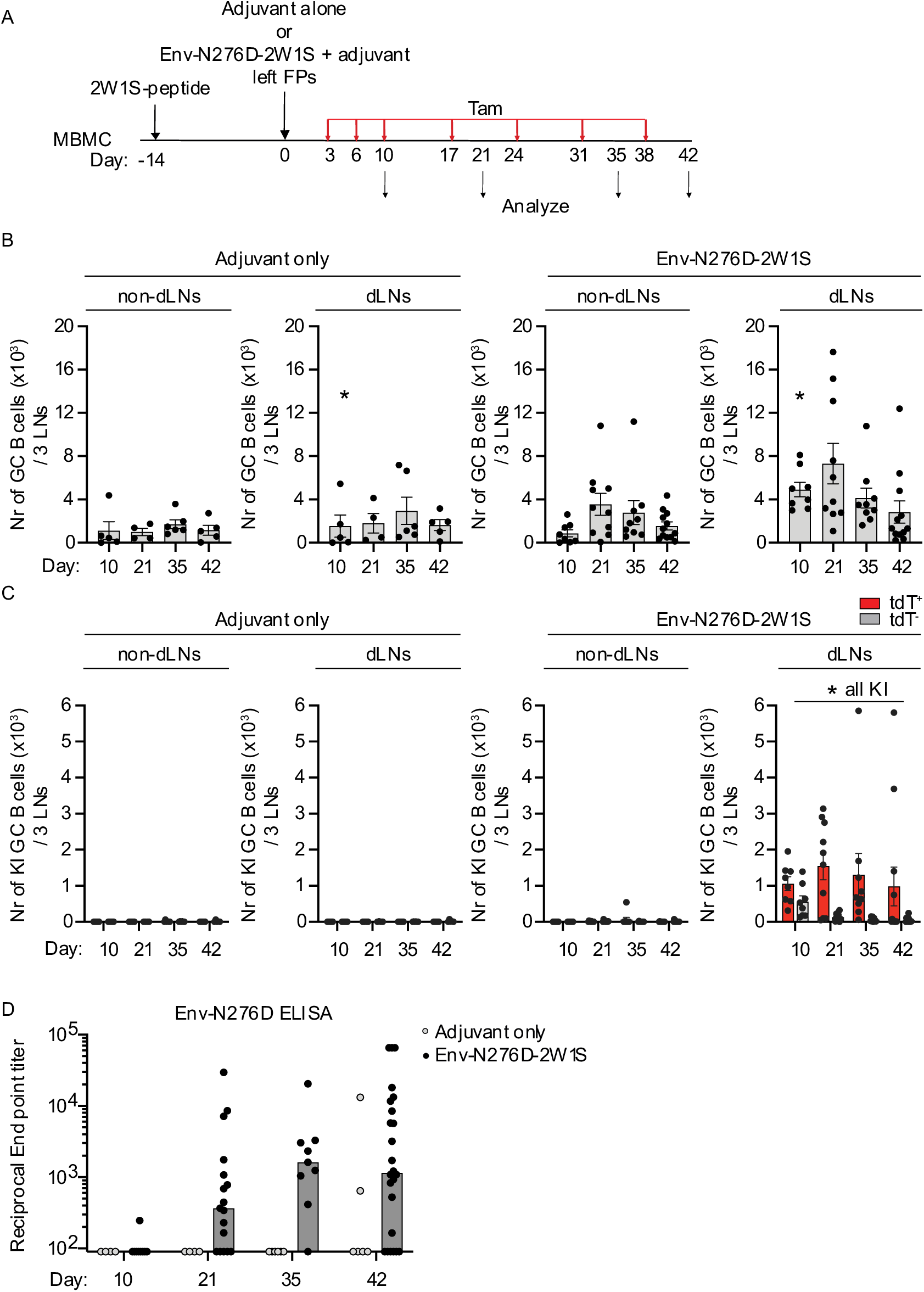
Immunization with intermediate affinity antigen and subsequent tamoxifen-treatment induces KI GC B cells of high labelling efficiency. (A) Schematic overview of the experiment: mixed bone marrow chimera mice were pre-primed intraperitoneally with 2W1S-peptide and immunized 2-4 weeks later in the left footpads with adjuvant only (control-immunized mice) or Env-N276D-2W1S in adjuvant. Mice were subsequently treated with tamoxifen at indicated timepoints (red arrows). Lymph nodes and serum antibodies were analyzed 10, 21, 35 and 42 days post immunization. (B) Graphs show number of GC B cells (GL7^+^, CD38^-^) per 3 LNs in non-draining and draining LNs over time in control- (adjuvant only) and Env-N276D-2W1S-immunized mice. (C) Graphs show number of KI GC B cells (GL7^+^, CD38^-^, CD45.2^+^) per 3 LNs in non-draining and draining LNs over time in control- (adjuvant only) and Env-N276D-2W1S-immunized mice. KI GC B cells are divided into fate-mapped in red (tdT^+^) and non-fate-mapped in gray (tdT^-^). (D) Graph shows N276D-specific IgG responses in serum over time from control- (adjuvant only, gray dots) and Env-N276D-2W1S- (black dots) immunized MBMC mice. Limit of detection is set at the lowest serum dilution (1/90). (B-D) Each dot represents one mouse and data is pooled from at least 2 independent experiments (total n= 4-29 per group). (B-C) Bars indicate mean and error bars show SEM. (D) Bars represent median. Statistical significance was determined using Mann-Whitney U test (*, P ≤ 0.05)

When we instead assessed the numbers and frequencies of KI GC B cells, we found very low to undetectable levels in both draining and non-draining LNs of control non-immunized mice. In Env-N276D-immunized mice, we detected KI GC B cells only locally in the draining LNs. Total numbers of KI GC B cells in draining LNs declined over time but were still above background at 42 days post immunization in 8 out of 12 mice (Figure 1C, Supplementary Figure 3A; Gating strategy 1, and Supplementary Figure 3C). The labelling efficiency of KI GC B cells with tdTomato was on average between 69-93%, depending on the time point post immunization (Supplementary Figure 3C and D).

Upon analysis of serum antibodies after immunization, we detected no significant binding to Env-N276D in control non-immunized mice (Figure 1D) confirming that the GC B cells detected in these mice were unspecific. However, mice that received Env-N276D showed detectable levels of Env-N276D-specific serum responses over time. In summary, these data confirm that intermediate affinity Env-N276D antigen induced an antigen-specific immune response in MBMC mice. KI GC B cells were efficiently fate-mapped upon tamoxifen-treatment and contributed to approximately 25% of total GC B cells in the draining LNs, with no significant occupancy in the non-draining LNs or in either LNs of control non-immunized mice.

### Somatically hypermutated GC- and memory KI B cells exhibit similar relative binding to Env-DMRS and wtEnv as the original 3BNC60 SI antibody

To determine the number of KI memory B cells elicited upon immunization, total fate-mapped KI B cells that formed after prime were analyzed based on their expression of GL7, CD38 and IgM (Supplementary Figure 3A; gating strategy 2). As shown in Figure 1C, fate-mapped KI GC B cells (tdT^+^, GL7^+^, CD38^-^) were present exclusively in draining LNs of immunized, but not control non-immunized mice, at all measured time points post immunization (Figure 2A). The number of IgM^+^ KI GC B cells decreased with time. When gating on KI memory B cells (tdT^+^, GL7^-^, CD38^+^) (Supplementary Figure 3A; gating strategy 2) we found the highest numbers (an average of 223 cells/3 LNs) in draining LNs at 10 days post immunization, which declined over time but were still detectable above background on day 42 (Figure 2B). Circulating KI memory B cells appeared in non-draining LNs 21 days post immunization and were also detectable above background on day 42. The majority of KI memory B cells in both draining- and non-draining LNs were IgM^+^ (Figure 2B). The amount of KI GC and memory B cells did not correlate with the frequency of naïve KI B cells prior to immunization (Supplementary Figure 4), indicating that a variation in KI B cell frequency at this level (an average of 1.8% of total B cells) did not affect the magnitude of the antigen-specific immune response by KI B cells after immunization as can be seen at lower frequencies of precursor cells ^48,57^. In summary, upon immunization with intermediate affinity Env-N276D immunogen, both switched and unswitched fate-mapped KI GC B cells formed in draining LNs of immunized mice. Fate-mapped KI memory B cells, the majority of which were IgM^+^, were initially detected only in draining LNs, but at later time points also in non-draining LNs, indicating the presence of circulating memory B cells.

**Figure 2:**
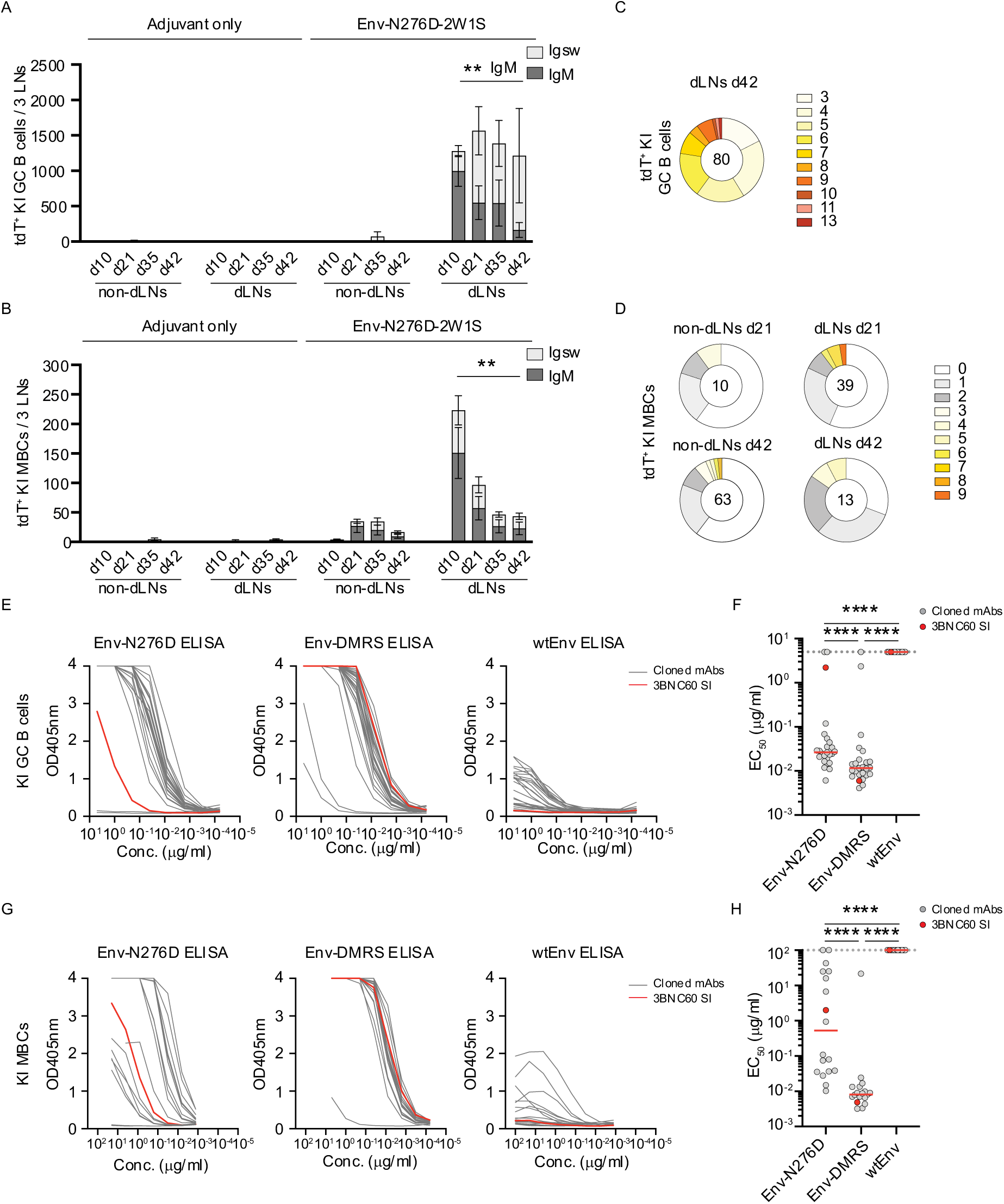
Somatically hypermutated GC- and memory KI B cells exhibit similar relative binding to wtEnv and Env-DMRS as the original 3BNC60 SI antibody. (A-B) Graphs show number of fate-mapped KI GC B cells (GL7^+^, CD38^-^, CD45.2^+^) (A) and fate-mapped KI memory B cells (MBCs) (GL7^-^, CD38^+^, CD45.2^+^) (B) per 3 LNs over time in control- (adjuvant only) and Env-N276D-2W1S immunized MBMC mice. ‘d’ indicates days post immunization. Isotype of the B cells are indicated in light gray (Igsw; switched) or dark gray (IgM). Data is pooled from 2 independent experiments (n=4-29 per group). Bars indicate mean and error bars show SEM. Statistical significance was tested using Mann-Whitney U test (**, P ≤ 0,01). (C-D) Pie charts depict the distribution of light chain nucleotide mutations of fate-mapped KI GC B cells sorted from ipsilateral LNs in MBMC mice 42 days after Env-N276D-2W1S-immunization (C) and fate-mapped KI memory B cells (MBCs) sorted from non-draining lymph nodes (non-dLNs) and draining lymph nodes (dLNs) in MBMC mice 21 and 42 days after Env-N276D-2W1S-immunization. (D) The number in the pie chart indicates the number of sequences analyzed. The slice size and colours indicate the fraction of sequences with the same number of nucleotide mutations. (E) ELISA data showing binding towards indicated Env antigens by the original 3BNC60 SI mAb (red) and mAbs cloned from somatically hypermutated fate-mapped KI GC B cells (gray) (n=24) indicated in (C). (F) Graph shows EC50 determined by ELISA for mAbs (gray) depicted in (E) towards Env-N276D, Env-DMRS and wtEnv. EC50 for the original 3BNC60 SI mAb is shown in red. Dotted line indicates limit of detection. (G) ELISA data showing binding towards indicated Env antigens by the original 3BNC60 SI mAb (red) and mAbs cloned from somatically hypermutated fate-mapped KI memory B cells (MBCs) (gray) (n=17) indicated in (D). (H) Graph shows EC50 determined by ELISA for mAbs (gray) depicted in (G) towards Env-N276D, Env-DMRS and wtEnv. EC50 for the original 3BNC60 SI mAb is shown in red. Dotted line indicates limit of detection. (F and H) Red line indicates median. Statistical significance was tested using Wilcoxon test (****, P ≤ 0,0001).

To investigate the effect of antigen affinity on the recall of fate-mapped KI GC B cells and KI memory B cells we aimed to boost mice that were primed with intermediate affinity Env (Env-N276D) with low or high affinity Env (wtEnv and Env-DMRS, respectively). Thus, we first verified if fate-mapped KI B cells still bound wtEnv and Env-DMRS with a relatively low and high affinity despite potential SHM diversification during the primary immune response. To do this, we single-cell sorted fate-mapped KI GC B cells present in draining LNs at day 42, as well as fate-mapped KI memory B cells from both draining- and non-draining LNs at day 21 and 42 post immunization. Upon sequencing the light chain (VJ) of the KI GC BCRs, we found that all sequences contained between 3 and 13 SHMs (Figure 2C). The KI memory B cells in draining LNs were less somatically hypermutated, with 44% of the cells containing 1 to 9 VJ SHMs at 21 days post immunization and 69% of the cells containing 1 to 5 VJ SHMs at 42 days post immunization (Figure 2D, left panels). Circulating KI memory B cells in non-draining LNs also expressed low levels of SHM, with 40% of the cells containing 1-4 and 1-8 LC SHMs, at 21 and 42 days after immunization, respectively (Figure 2D, right panels).

We produced monoclonal antibodies of paired heavy- and light antibody chains representing selected BCRs from sequenced fate-mapped KI GC B cells (Supplementary Figure 5) and all BCRs from sequenced fate-mapped KI memory B cells with non-synonymous mutations. When first tested in ELISA against Env-N276D (the immunogen the mice were initially primed with), we found that 22/24 cloned KI GC BCRs bound with a lower EC50 value compared to the original 3BNC60 SI mAb. Compared to 3BNC60 SI, all but two KI GC BCRs bound with a similar EC50 value to Env-DMRS. 3BNC60 SI does not bind detectably to wtEnv in ELISA and even though a proportion of the cloned KI GC BCRs displayed detectable binding, it was too low to calculate an accurate EC50 (Figure 2E). If the EC50 values of the cloned KI GC BCRs to wtEnv were approximated to the lowest level of detection, we found that the GC BCRs bound Env-DMRS with at least a 16-fold lower EC50 compared to wtEnv (Figure 2F). For the cloned KI memory BCRs, 10/17 bound with a lower EC50 to Env-N276D compared to the original 3BNC60 SI. All but one KI memory BCR bound with a similar EC50 value to Env-DMRS as 3BNC60 SI and 9/17 KI memory BCR displayed detectable binding to wtEnv (Figure 2G). If the EC50 values of the cloned KI memory BCRs to wtEnv were approximated to the lowest level of detection, KI memory BCRs bound Env-DMRS with a 96-fold lower EC50 value compared to wtEnv (Figure 2H). We conclude that the fate-mapped GC and memory KI B cells harboring SHM still bound significantly better to Env-DMRS than to wtEnv. Importantly, this confirms that wtEnv and Env-DMRS can be used to evaluate the role of antigen affinity on GC B cell and memory B cell recall responses in mice primed with Env-N276D antigen.

To study recall responses we primed MBMCs with intermediate affinity Env-N276D in the left foot pads. Mice were treated with tamoxifen to label GC B cells and then boosted at 42 days post prime in all four foot pads with either low affinity wtEnv or high affinity Env-DMRS (Figure 3A). Since tamoxifen is only given after prime, naïve B cells that get *de novo* activated upon boost will not be fate-mapped.

**Figure 3:**
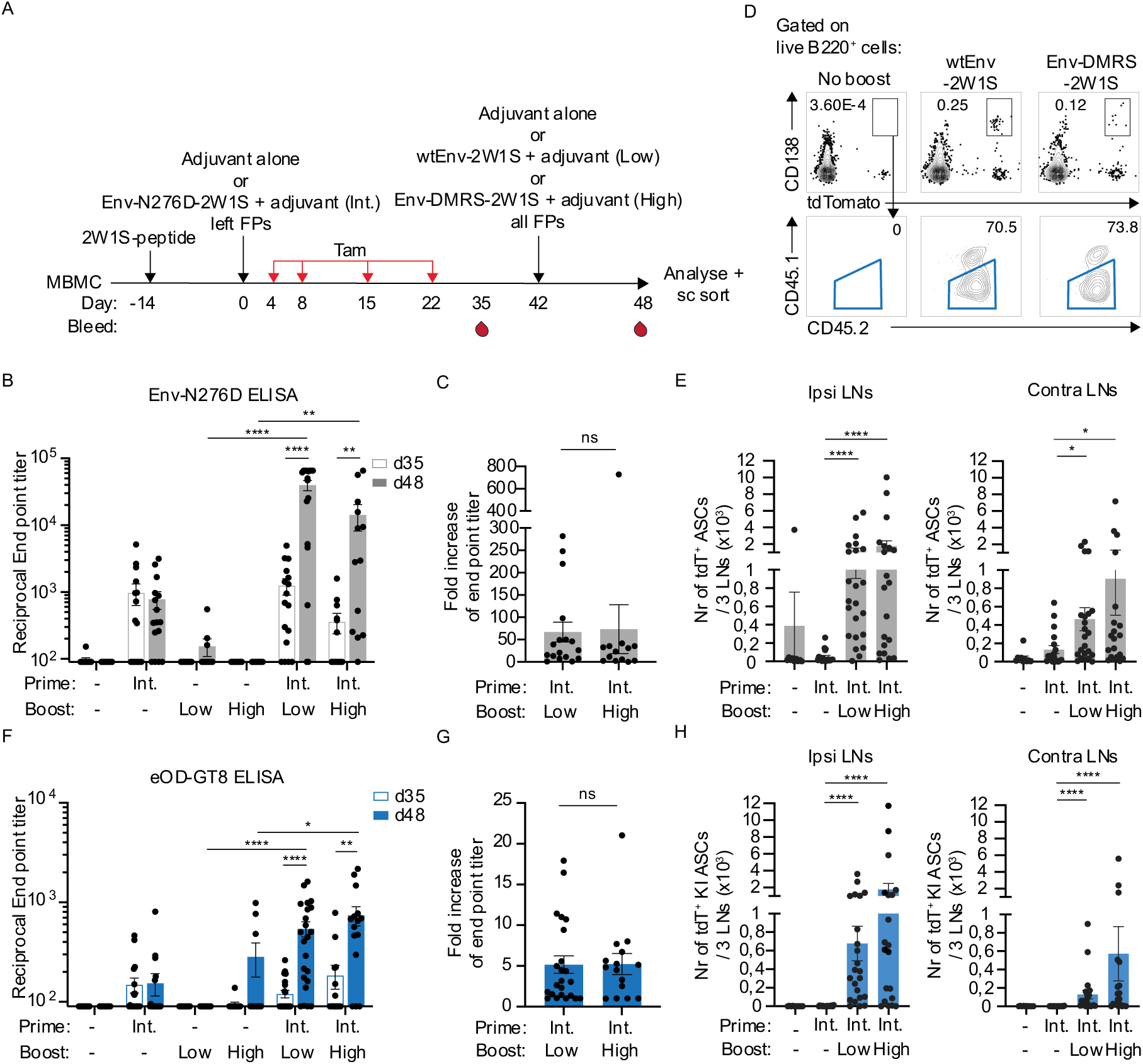
Levels of pre-existing antibodies elicited by Env-N276D does not interfere with serum recall responses of KI memory B cells. (A) Schematic overview of the experiment: mixed bone marrow chimera (MBMC) mice were pre-primed intraperitoneally with 2W1S-peptide and immunized 2-4 weeks later in the left footpads (FPs) with adjuvant only (=control-immunized mice) or intermediate affinity (Int.) Env-N276D-2W1S in adjuvant. ‘-‘ indicates immunization with adjuvant only. Mice were subsequently treated with tamoxifen (Tam) at indicated timepoints (red arrows) and boosted 42 days post prime in all 4 footpads with either adjuvant only, low affinity wtEnv-2W1S (Low) or high affinity Env-DMRS-2W1S (High) in adjuvant. Analysis of serum was performed at day 35 (7 days pre boost) and 48 (6 days post boost) and analysis of lymphocytes was performed on day 48. (B) Graph shows serum IgG responses against Env-N276D of MBMC mice treated as in (A) at day 35 (pre-boost, white bars) and 48 (post-boost, gray bars). ‘d’ indicates days after prime. Limit of detection is set at the lowest serum dilution (1/90). (C) Graph shows fold-increase in reciprocal end point titer of serum antibody response in (B) between day 6 post-boost and day 35 post-prime. (D) Representative flow cytometry plots from the experiment in (A) showing the frequency of fate-mapped antibody secreting cells (ASCs) (upper panels) and fate-mapped KI ASCs (lower panels) 48 days after prime only (no boost), or 6 days post-boost with wtEnv-2W1S or Env-DMRS-2W1S. (E) Graphs show the number of fate-mapped ASCs per 3 ipsi- or contralateral LNs in MBMC mice at day 48, when treated as in (A). (F-G) As in (B-C) but against eOD-GT8. (H) Graphs show the number of fate-mapped KI ASCs per 3 ipsi- or contralateral LNs in MBMC mice at day 48, when treated as in (A). (B-C and E-H) Each dot represents one mouse and data is pooled from at least 3 independent experiments (n=10-23 per group). Bars indicate mean and error bars show SEM. Statistical significance was tested using Mann-Whitney U test (*, P ≤ 0.05; **, P ≤ 0.01;****, P ≤ 0.0001; ns = non-significant).

### Levels of pre-existing antibodies elicited by Env-N276D does not interfere with serum recall responses of KI memory B cells

High levels of pre-existing antibodies generated upon prime can interfere with recall responses upon secondary immunization ^39,42–47^. Therefore, we first evaluated if memory B cells generated after prime in our mouse model are successfully activated to become antibody-secreting cells upon boost. To first determine the level of serum antibodies pre- and post-boost, we performed an ELISA against Env-N276D of serum collected 35 days post prime and 6 days post boost (d48). As shown before on day 35 post prime, Env-N276D-specific serum responses were present in Env-N276D-immunized, but not in control non-immunized mice (Figure 3B). After boosting with either wtEnv or Env-DMRS, we detected a significant increase in Env-N276D-specific serum responses for both antigens. The antibody levels post boost were higher than in mice that only received a single immunization on day 42 with no previous prime, indicating differentiation of memory B cells into antibody-secreting cells upon boost (Figure 3B). The fold change in serum antibody end point titers between pre- and post-boost, were similar with both low and high affinity antigen (Figure 3C), indicating that the overall boost effect on the serum antibody level was not significantly affected by antigen affinity. We also found similarly increased levels of fate-mapped antibody-secreting cells by flow cytometry (tdT^+^, CD138^+^) in both the LNs contralateral (contra; right) and ipsilateral (ipsi; left) to the side of the initial prime (left) 6 days after both low and high affinity antigen boost (Figure 3D and E).

To specifically determine the serum antibody response generated by KI B cells, we made use of the differential binding properties of the KI Abs to another Env-based immunogen, eOD-GT8. Due to the heterogeneity between the HxB2-based eOD-GT8 and the 426c-based Env immunogens, polyclonal non-CD4bs responses elicited by Env-N276D are not detected in the eOD-GT8-ELISA while 3BNC60 SI binding to the conserved CD4bs is retained. This was confirmed by analyzing the serum of Env-N276D-immunized WT or MBMC mice in ELISA where serum from WT mice only responded to Env-N276D whereas serum from MBMC mice responded to both Env-N276D and eOD-GT8 (Supplementary Figure 6).

eOD-GT8-specific serum antibodies were detected in immunized, but not control non-immunized mice (Figure 3F). Similar to Env-N276D-specific serum antibodies post boost, we detected a significant increase in eOD-GT8-specific serum antibodies upon boost with both antigens which were higher than in mice that only received a single immunization on day 42 with no previous prime (Figure 3F). The fold change in serum antibody end point titers between pre- and post-boost against eOD-GT8 was similar with both antigens (Figure 3G). Moreover, we detected increased levels of KI antibody-secreting cells (tdT^+^, CD45.2^+^, CD138^+^) in both contra- and ipsilateral LNs in mice after both low and high affinity antigen boost but not in control non-immunized or non-boosted mice (Figure 3 D and H). Overall, this data shows that the level of pre-existing antibodies in our mouse model at day 42 post prime, is not blocking the recall of fate-mapped B cells to form antibody-secreting cells. Furthermore, we found that there was no detectable difference in the recall response on a serum level induced by the different affinity antigens.

### High affinity antigen boost results in higher levels of memory-derived GC B cells than low affinity antigen boost

To evaluate the effect of antigen affinity for memory B cell entry into secondary GCs upon boost, we analyzed the LNs contralateral to the primed side on day 6 post boost of the experiment outlined in Figure 3A by flow cytometry (Figure 4A). Importantly, since no fate-mapped KI GC B cells were detected after prime in the LNs of the contralateral site (Figure 2A), if present after boost, they can be defined as memory-derived GC B cells. In contrast, non-fate-mapped KI GC B cells in secondary GCs are most probably derived from naïve B cells ^34,47^.

**Figure 4:**
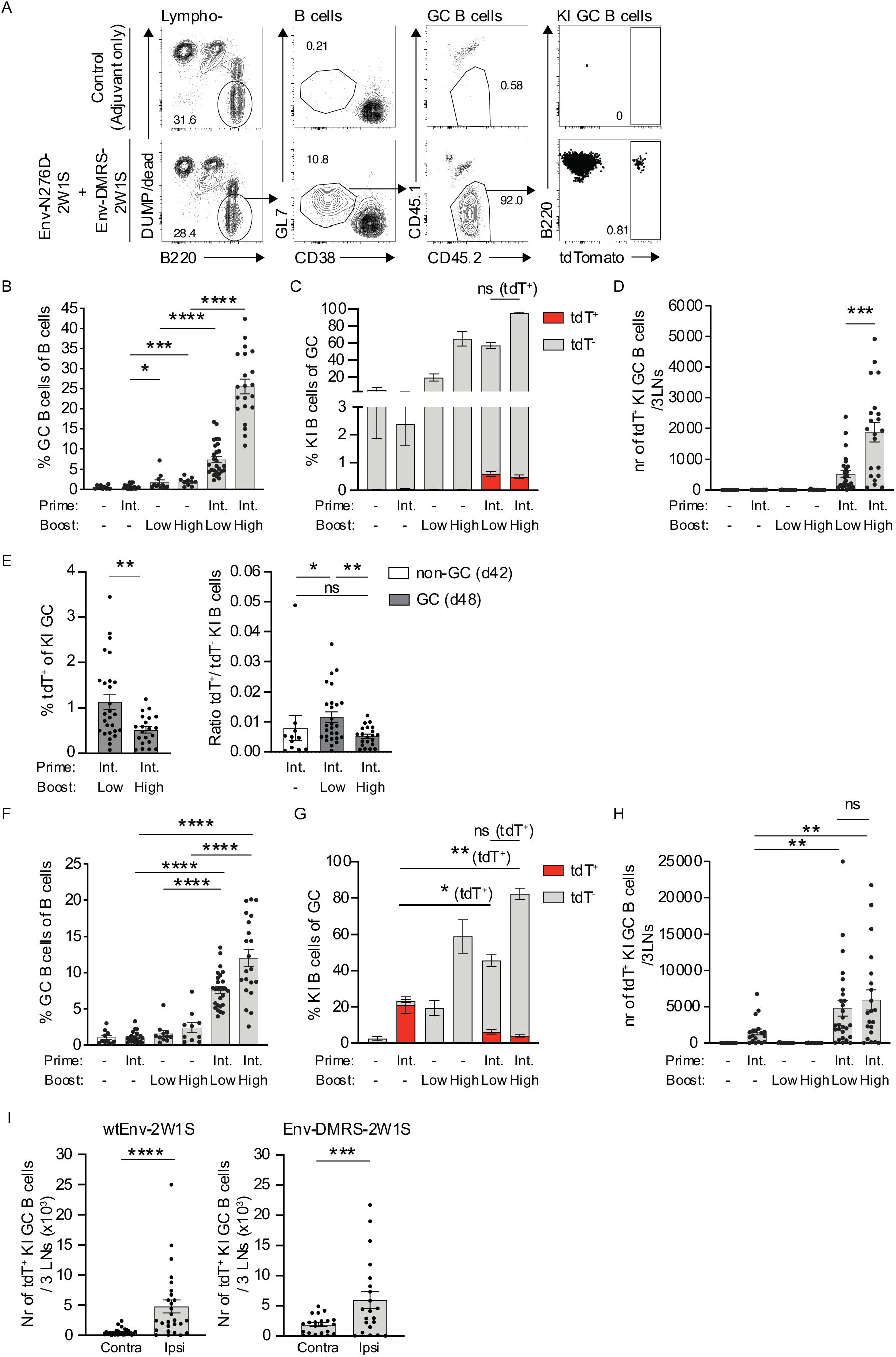
Antigen affinity affects the number of previously activated B cells present in secondary GCs in the contralateral but not the ipsilateral side. Analysis of MBMC mice treated as outlined in Figure 3A. (A) Representative flow cytometry plots showing gating for GC B cells (GL7^+^, CD38^-^), KI GC B cells (CD45.2^+^) and fate-mapped (tdT^+^) KI GC B cells in the contralateral lymph nodes of MBMC mice at 6 days post boost. (B-D) Graphs show frequencies GC B cells (GL7^+^, CD38^-^) of B cells in contralateral lymph nodes (B), frequencies KI B cells (CD45.2^+^) of GC B cells in contralateral lymph nodes (C) and number of fate-mapped (tdT^+^) KI GC B cells per 3 contralateral lymph nodes (LNs) (D) at day 48. (E) Frequency fate-mapped (tdT^+^) KI GC B cells of KI GC B cells (left panel) and ratio of fate-mapped (tdT^+^) KI B cells over non-fate-mapped (tdT^-^) KI B cells at day 42 in the non-GC compartment (white bar) and at day 48 in the GC compartment (gray bars) in the contralateral side 6 days post boost (right panel). (F-H) Graphs show frequencies GC B cells of B cells in ipsilateral lymph nodes (F), frequencies KI B cells (CD45.2^+^) of GC B cells in ipsilateral lymph nodes (G) and number of fate-mapped (tdT^+^) KI GC B cells per 3 ipsilateral lymph nodes (LNs) (H) at day 48. (I) Graphs show total number of fate-mapped (tdT^+^) KI GC B cells per 3 lymph nodes (LNs) in contra-(Contra) versus ipsilateral (Ipsi) LNs after wtEnv-2W1S boost (left panel) or Env-DMRS-2W1S boost (right panel). (C and G) Bars represent stacked non-fate-mapped (tdT^-^; gray) and fate-mapped (tdT^+^; red) KI B cell frequencies of GC B cells. (B-I) Each dot represents one animal and data is pooled from at least 3 independent experiments (n= 10-27 per group). ‘Int.’ indicates ‘intermediate affinity antigen’ Env-N276D-2W1S, ‘Low’ indicates ‘low affinity antigen’ wtEnv-2W1S, ‘High’ indicates ‘high affinity antigen’ Env-DMRS-2W1S and ‘-‘ indicates immunization with adjuvant only. Bars indicate mean and error bars show SEM. Statistical significance was tested using Mann-Whitney U test or Wilcoxon test (*, P ≤ 0.05; **, P ≤ 0.01; ***, P ≤ 0.001; ****, P ≤ 0.0001; ns = non-significant).

In contralateral LNs of control non-immunized mice or mice that were only primed, we detected low background GC B cells (0.6% of total B cells) (Figure 4B) that consisted almost exclusively of polyclonal B cells (Figure 4C). In mice that were only given one immunization on day 42, we found higher GC B cell frequencies, of which an average of 19 and 65% were KI B cells after immunization with low and high affinity antigen, respectively. As expected, since tamoxifen was only administered between day 4 and 22, all these cells were non-fate-mapped (Figure 4B and C). In contrast, after boosting mice that were previously primed, we found that low and high affinity antigen induced increased GCs (7.0 and 26%, respectively) (Figure 4B) constituting 57 and 95% KI B cells, respectively (Figure 4C). The differently sized GCs post-boost are most probably due to an affinity-dependent influx of naïve 3BNC60 SI KI B cells ^48^. Importantly, only an average of 0.59% and 0.49% of GC B cells were KI memory B cell-derived (fate-mapped) after boost with either low or high affinity antigen, respectively (Figure 4C). This correlates well with previous findings in mice, that mainly naïve B cells participate in secondary GCs upon boost ^34,40^. However, even though the proportion of memory-derived KI B cells of total GC B cells were similar between the different affinity antigens upon boost, the absolute numbers were 3.6–fold higher after boosting with the high affinity antigen, compared to the low affinity antigen (Figure 4D). Similar results were detected in repeat experiments where the boost was only given contralateral to the prime, indicating that the simultaneous boost-immunization in the ipsilateral side does not affect the events in the contralateral LNs (Supplementary figure 7A-D).

Interestingly, when excluding the polyclonal GC B cells and focusing on the KI B cell compartment of the GC in contralateral LNs, the proportion of memory-derived B cells was higher when using low affinity, compared to high affinity antigen boost (Figure 4E, left panel). To determine if memory B cells showed an advantage over naïve B cells to enter and/or expand in secondary GCs, we analyzed the ratio of fate-mapped over non-fate-mapped KI B cells in the non-GC compartment (GL7^-^, CD38^+^) pre-boost and in the GC-compartment (GL7^+^, CD38^-^) post-boost. Compared to the non-GC compartment pre-boost, the ratio was higher in the GC compartment after boosting with wtEnv, but not with Env-DMRS (Figure 4E, right panel). In summary, our data suggests that a high affinity antigen results in more memory-derived GC B cells in secondary GCs compared to a low affinity antigen. Moreover, memory B cells exhibited an advantage over naive B cells to enter and/or expand in secondary GCs upon low, but not high, affinity boost.

### Refueling ongoing GCs leads to an accumulation of fate-mapped GC B cells in an antigen affinity-independent mode

A second immunization in the local, draining site, where residual GCs are still present from a first immunization, was referred to as ‘refueling’ of a GC ^34^. Similar to when performing a ‘long prime’, using osmotic pumps or performing repetitive immunizations within a short time window ^58–60^, this could lead to sustained or increased GC B cell proliferation and thereby increased levels of SHMs in GC B cells that were initiated upon prime. GC refueling will however most probably also lead to an influx of activated naïve or memory B cells ^40,61,62^ that could potentially affect residual GC B cells to leave GCs ahead of time. In addition, it has previously been shown that refueling ongoing GCs with soluble antigen may lead to increased GC B cell death though apoptosis ^63,64^.

To investigate the process of refueling in more detail, we analyzed the LNs ipsilateral to the primed side on day 6 post boost in the experiment outlined in Figure 3A. In mice primed and boosted with only adjuvant, we detected low levels of GC B cells (an average of 1.0% of B cells) containing exclusively polyclonal B cells (Figure 4F and G). Primed mice that were not boosted showed low levels of residual GCs (an average of 1.0% of total B cells) containing an average of 22% KI B cells of which the majority was fate-mapped and hence represented residual KI GC B cells from the prime immunization. In control mice that were only given one immunization at day 42 with either low- or high affinity antigen, we detected GC B cells (an average of 1.6 and 2.4% of total B cells, respectively) of which a large proportion were KI B cells (an average of 19 and 59% of total GC B cells, respectively). As expected, since tamoxifen was only administered between day 4 and 22, none of these KI GC B cells were fate-mapped (Figure 4F and G). In contrast, in mice that were previously primed and then received a boost, we found that low and high affinity antigen induced increased levels of GC B cells (an average of 7.6 and 12% of total B cells, respectively) compared to mice that only received one immunization on day 0 or day 42 (Figure 4F). These GCs consisted of an average of 46 and 82% KI B cells after low and high affinity antigen boost, respectively. Compared to mice that were primed but not boosted, the proportion of fate-mapped KI GC B cells was lower, and similar for both boosting antigens (an average of 6.2 and 4.1% of total GC B cells for low and high affinity antigen respectively) (Figure 4G). Importantly however, when we instead analyzed the total numbers of fate-mapped KI GC B cells, we found a 3-4-fold increase in boosted, compared to non-boosted mice and this increase was independent of antigen affinity (Figure 4H).

When compared to contralateral LNs after boost, the numbers of fate-mapped KI GC B cells in the ipsilateral LNs were nine-fold and three-fold higher for low and high affinity antigen, respectively (Figure 4I). In conclusion, our data show that boosting the same site as previously primed results in a refueling effect with increased numbers of fate-mapped KI GC B cells. This increase was independent of boosting antigen affinity and could be the combined result from both fate-mapped memory B cell entry to refueled GCs and an expansion of residual GC B cells, originally activated at prime.

### Low affinity boost results in recalled B cells in secondary GCs with higher levels of SHM compared to high affinity boost

To gain further qualitative insight into the memory-derived KI GC B cells in the contralateral LNs post boost, we performed SHM analysis of 266 and 204 single-cell sorted fate-mapped and 108 non-fate-mapped KI GC B cells after either low or high affinity antigen boost, respectively (Figure 5A, Supplementary figure 8A). Approximately 30% of the non-fate-mapped KI GC B cells (naïve B cell-derived) were somatically hypermutated with an average of 1.6 SHMs, (Figure 5A, Supplementary Figure 8B). This indicates that SHM levels above this in fate-mapped cells would reflect SHMs that were present already upon entry into the secondary GC. When comparing the SHM of non-fate-mapped and fate-mapped GC B cells in secondary GCs, we detected significantly higher numbers in the fate-mapped cells after low affinity boost, but not after high affinity boost. When directly comparing the fate-mapped cells, the low affinity boosted cells showed higher levels of SHMs compared to the high affinity boosted cells. Furthermore, the numbers of SHMs in fate-mapped GC B cells after low affinity, but not after high affinity boost, was higher when compared to the SHMs of circulating memory B cells present in the contralateral LNs on the day of the boost (Figure 5A). Non-synonymous mutations of fate-mapped GC B cells were distributed over the whole VJ sequence for both boosting antigens, with a higher mutation frequency associated to the CDR regions (Figure 5B). Not all fate-mapped B cells contained SHMs and when comparing the fraction of somatically unmutated memory B cells pre-boost, with the memory B cells that entered secondary GCs, we found that the frequency of sequences that expressed no SHM decreased from 60 to 36% after low affinity antigen boost but stayed approximately equal (62%) after high affinity boost (Figure 5A). These results were similar when repeated in mice that were only boosted in the contralateral LNs, indicating that the simultaneous boost-immunization in the ipsilateral site does not affect the overall events in the contralateral LNs (Supplementary figure 8C and D). This data shows that the lower affinity antigen selectively recruits memory B cells into secondary GCs with higher levels of SHM upon entry compared to the higher affinity antigen.

**Figure 5:**
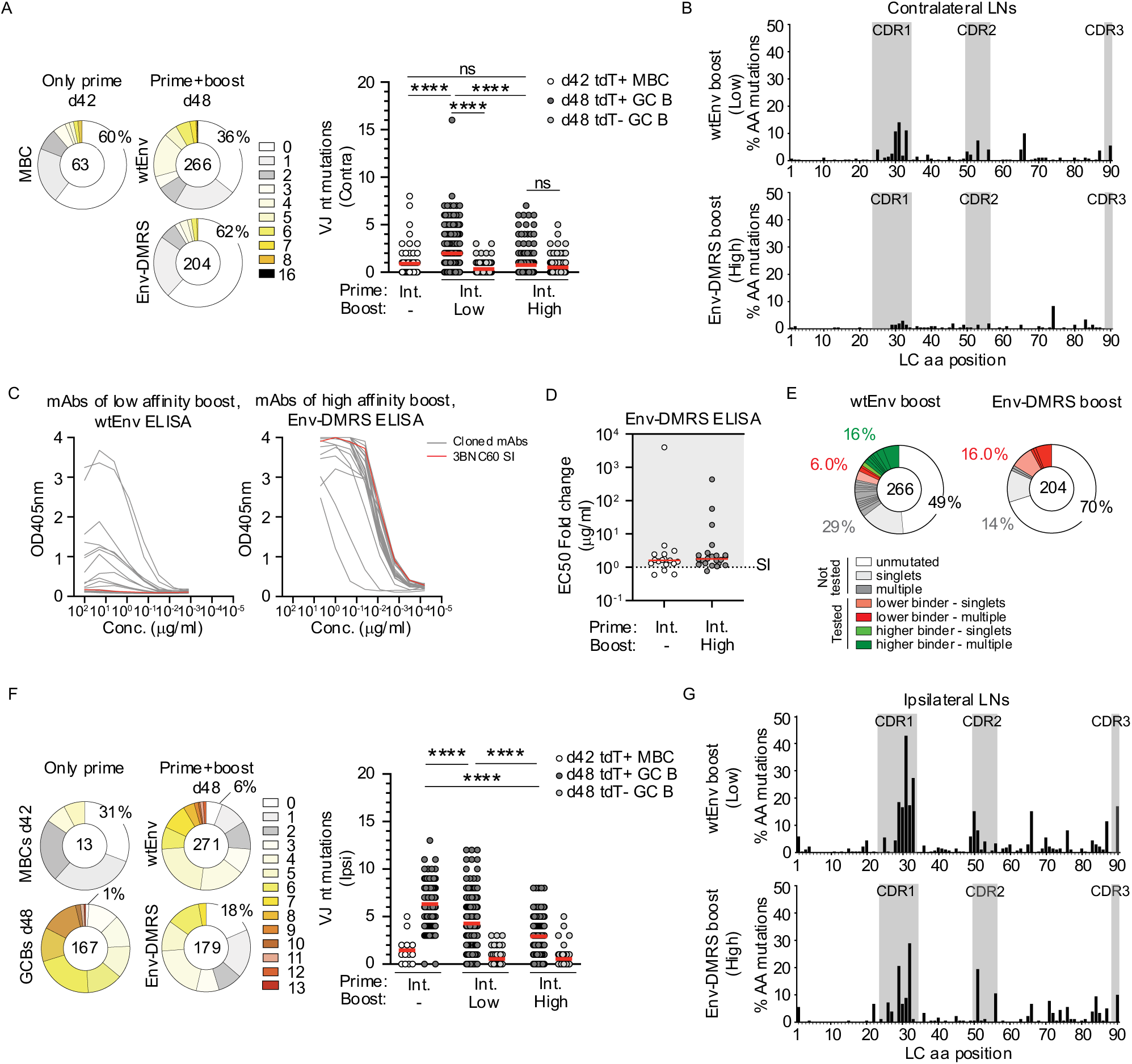
Low affinity boost results in recalled B cells in secondary GCs with higher levels of SHM compared to high affinity boost. (A) Pie charts depict the frequency of light chain (VJ) nucleotide mutations of fate-mapped KI memory B cells (left) and fate-mapped KI GC B cells (right) single cell sorted from contralateral LNs 42 days after Env-N276D-2W1S immunization and 6 days after wtEnv-2W1S or Env-DMRS-2W1S boost, respectively. The number in the pie chart indicates the number of sequences analyzed. The slice size and colours indicate the fraction of sequences with the same number of nucleotide mutations. The percentage indicate frequency of unmutated sequences. Graph shows the number of light chain (VJ) nucleotide mutations of fate-mapped KI memory B cells (white) single-cell sorted from contralateral LNs 42 days after Env-N276D-2W1S immunization, fate-mapped (tdT^+^; dark grey) and non fate-mapped (tdT^-^; light gray) KI GC B cells single-cell sorted from contralateral lymph nodes 6 days after wtEnv-2W1S or Env-DMRS-2W1S boost. (B) Graphs shows the distribution of amino acid mutations in the light chain sequences of fate-mapped KI GC B cells single cell sorted from contralateral lymph nodes 6 days after wtEnv-2W1S (upper panel) or Env-DMRS-2W1S (lower panel) boost. (C) ELISA data showing binding towards indicated Env antigens by the original 3BNC60 SI mAb (red) and mAbs cloned from somatically hypermutated fate-mapped KI GC B cells (gray) (n=43) indicated in (A). (D) Graph shows EC50 determined by ELISA for mAbs cloned from fate-mapped KI memory B cells (white) depicted in (A) and fate-mapped KI GC B cells (gray) depicted in (C) towards Env-DMRS-2W1S. EC50 for the original 3BNC60 SI mAb is indicated by the dotted line. (E) Pie charts depict the distribution of light chains of fate-mapped KI GC B cells as in (A). Slice colour correlate to sequences with no amino acid mutations (white) and sequences with amino acid mutations that were not cloned and tested (gray) or sequences with amino acid mutations that were cloned and tested for binding in ELISA in (C) (green = ‘higher binders’, red = ‘lower/similar binders’). Sequences were divided based on number of identical sequences found (light colour = singlets, darker colour = multiple). Percentages show frequency of sequences in the corresponding pie slice. (F-G) As in (A-B) but for ipsilateral LNs. (A-B and F-G) Data is pooled from at least 3 independent experiments with 4-6 mice per group. Each dot represents the VJ sequence from one cell (n= 13-271 per group). Red line indicates mean values. Statistical significance was tested using Mann-Whitney U test (****, P ≤ 0.0001; ns = non-significant).

So far, upon comparing low and high affinity boosting antigens, we made three observations that were unique to the low affinity boost immunization; First, low affinity boosting antigen preferentially stimulated memory KI B cells over naïve KI B cells to enter secondary GCs (Figure 4E). Second, low affinity boosting antigen recruited affinity-matured memory B cells into secondary GCs with higher levels of SHM upon entry, and third, compared to the circulating memory B cell pool present pre boost, the proportion of memory B cell-derived cells in secondary GCs with 0 SHM are lower (Figure 5A). Together, these findings suggest that there is an affinity threshold for memory B cells to enter secondary GCs upon recall. To evaluate this further, we produced 43 monoclonal antibodies representing selected BCRs with non-synonymous mutations of fate-mapped B cells from secondary GCs in contralateral LNs after low- (22) and high (22) affinity antigen boost (Supplementary Figure 9A and B). When testing the 22 monoclonal antibodies of memory B cell-derived GC B cells induced by the low affinity antigen in a wtEnv-ELISA, we found that 11 of the 22 cloned antibodies bound with an increased binding phenotype (‘higher binders’) compared to the original 3BNC60 SI antibody, expressed by the naïve KI B cells (Figure 5C). In contrast, none of the 22 cloned antibodies of memory B cell-derived GC B cells induced by the high affinity antigen showed increased binding to Env-DMRS in ELISA compared to 3BNC60 SI (Figure 5C and D). Importantly, the ‘higher binders’ detected after low affinity antigen boost, more often represented sequences that were found multiple times in the fate-mapped B cells (Figure 5E). Collectively, these results support the idea that an affinity threshold for memory B cell entry to secondary GCs exist upon boost, where memory B cells that bind more potently to the low affinity antigen are favored in the GC-entry process over the cells expressing lower affinity BCRs.

To understand the process of refueling in more detail, we also performed SHM analysis of 271 and 179 single cell sorted fate-mapped KI GC B cells in the ipsilateral LNs after boosting with either low or high affinity antigen, respectively (Figure 5F, Supplementary figure 8E). We unexpectedly found that even though the fate-mapped population had increased in total numbers independent of the boosting antigen affinity (Figure 4H), when compared to fate-mapped GC B cells of ongoing GCs at day 48 that had not been refueled, the average levels of SHM were significantly lower after boosting with either antigen. In agreement with what we found in the contralateral side, the level of SHM was the lowest with the high affinity antigen (Figure 5F, Supplementary figure 8E). The number of fate-mapped cells with no SHM increased from 1% in the ongoing GCs that were not refueled, to 6% after boost with the low-and 18% after boost with the high affinity antigen. For the high affinity boosting antigen, in addition to detecting an accumulation of fate-mapped cells with low levels of SHM, we also detected a loss of cells with high numbers of mutations (9-13 SHMs), present in the ongoing GCs that were not refueled (Figure 5F). The SHMs of fate-mapped GC B cells were distributed over the whole VJ sequence for both boosting antigens, with a higher mutation frequency associated to the CDR regions (Figure 5G). In summary, even though the total number of fate-mapped cells increased similarly after both low and high affinity antigen boost in the ipsilateral LNs, the composition of these cells post boost differed depending on the antigen affinity. The decreased levels of SHMs, which was more pronounced with high affinity antigen, indicate that a large proportion of fate-mapped cells post boost is a result of memory B cells that enter GCs, similar to what we detected after boost in contralateral LNs, rather than the expansion of residual fate-mapped GC B cells. Furthermore, we reason that the loss of B cell clones present in ongoing GCs with the highest levels of SHM upon high affinity, but not low affinity boost, suggest a maximum affinity threshold of the GC B cells, where the GC B cell clones of highest affinity exit the GC upon antigen stimulation.

## Discussion

To design novel prime/boost vaccines that protect against genetically variable pathogens or pathogens that use other strategies to evade the immune response, we need a more detailed understanding of the regulation of B cell recall responses, and in particular, how memory B cells enter GCs. Here, using a mouse model with both polyclonal-, and HIV-1 Env-specific KI B cells that express fate-mapping genes and bind to HIV-1 Env-proteins with specific affinities, we studied the cellular effects of antigen affinity for B cell recall responses. We found that a relatively high affinity boosting antigen resulted in increased numbers of memory-derived B cells in secondary GCs. Furthermore, our results showed that a low affinity boosting antigen more selectively promoted memory B cells with higher levels of SHM to enter secondary GCs. Taken together, these findings suggest an affinity threshold for memory B cell GC entry. Moreover, we found that refueling local LNs leads to increased numbers of GC B cells originally activated by prime, but with significantly lower average levels of SHM compared to non-boosted draining LNs, indicating high levels of memory B cell entry upon refueling.

In mouse models, the entry of memory B cells into secondary GCs is a rare event ^34,40^. Memory B cells with specific phenotypes are more associated with GC entry than others, including the expression of low levels of CD80 and PDL2 ^36^ and the expression of the IgM isotype, even though IgG memory B cells are also capable of GC entry ^37,38^. By boosting and studying secondary GCs in LNs contralateral to the primary immunization, upon fate-mapping of primary GCs, we could distinguish between B cells in secondary GCs that were memory B cell-derived (fate-mapped by tdTomato) and B cells that had not participated in earlier GCs (non-fate-mapped by tdTomato). As was shown previously using similar immunization strategies ^34,40^, we found that secondary GCs consisted mainly of naïve B cells that were *de novo* activated upon boost and only a minority of the GC B cells were memory-derived. Non-fate-mapped cells in secondary GCs could also originate from non-GC dependent, early memory B cells generated upon prime that were not labeled by tdTomato due to the low expression of S1PR2 ^56^. However, recent experiments by Schiepers *et al.* using a mouse model that constitutively labeled all naïve B cells present prior initial immunization, suggests that the majority of B cells in secondary GCs after boost are unlabeled, indeed confirming a naïve, rather than an early memory B cell origin of the majority of cells in secondary GCs after boost ^47^.

The effect of pre-existing antibodies in a sequential vaccination strategy has been discussed intensively since it was shown that antibodies could block or modify the activation of B cells with the same specificity upon immunization ^39,42,45,47^. It is interesting to note that the limited entry of memory B cells into secondary GCs in our system, was most probably not due to competition for the antigen with pre-existing cognate antibodies generated during prime, since naïve KI B cells of the same specificity (non-fate-mapped), were successfully recruited and represented most B cells in the secondary GCs.

The proportion of memory-derived GC B cells in secondary GCs were similarly low with high- and low affinity antigen boost. However, since we are using a MBMC mouse model with increased levels of naïve KI B cells that can be activated upon boost, this resulted in significantly larger GCs when boosting primed mice with the high-, compared to the low affinity antigen. To deconvolute the data, we instead counted the absolute numbers of fate-mapped memory-derived GC B cells and found higher numbers when using the high affinity antigen. Importantly, these cells displayed lower levels of SHM compared to when boosting with low affinity antigen. We reasoned that the increased numbers of memory-derived GC B cells after high affinity antigen boost could either be due to the recruitment of more memory B cells to the GCs, or alternatively, the memory B cells that entered the GC divided more compared to when boosting with the low affinity antigen. The overall low levels of SHMs in these cells and the fact that the SHMs were not extensively shared between the VJ sequences would however suggest that more memory B cells were recruited with a high affinity antigen. In contrast, the higher level of SHM in the fate-mapped memory-derived GC B cells after low affinity boost could result from more division in the secondary GC, however, the fact that non-fate-mapped KI B cells present in the secondary GC at the same time point showed significantly lower SHM, indicates that a higher level of SHM was present in the fate-mapped B cells already upon GC entry.

We furthermore found that compared to the high affinity antigen boost, with a low affinity boost, memory B cells showed a selective advantage over naïve B cells with the same specificity to enter secondary GCs and a reduced fraction (36% compared to 62% with high affinity antigen) of the GC-recruited memory B cells had no SHM. Upon cloning BCRs from memory-derived GC B cells after low affinity boost, we found that they bound the boosting antigen better than the original mAb 3BNC60 SI to a higher degree than the memory-derived GC B cells after high affinity boost. Taken together, our data studying the entry of memory B cells to secondary GCs after different affinity antigen boost, suggests that there is a selection for memory B cell GC entry based on the affinity to the antigen, similar to what was shown previously for naïve B cell GC entry ^65^. The presence of an affinity-threshold for memory B cell entry to GCs also agrees with previous findings in another HIV-1 antibody KI mouse model where a sequential immunization approach using lower affinity antigens as boost resulted in KI BCRs with higher levels of SHM compared to when repeatedly immunizing with a high affinity antigen ^21^.

Prime boost vaccine regimens are often given locally in the same immunization site, and the result is evaluated by determining serum antibody responses, which increase after boost due to the differentiation of memory B cells into antibody-producing cells ^32^. The cellular events that take place in draining LNs during boost has however not been clearly investigated. By studying GCs in LNs after boosting ipsilateral to the primary immunization, after fate-mapping B cells participating in the primary GCs, we could examine the effects of refueling residual GCs with different affinity antigens. In contrast to the contralateral side, the fate-mapped B cell population after boost could here be a mixture of GC B cells activated upon prime that never left the GC and stayed in the GC after boost, and memory B cells generated by the prime that entered GCs in response to the boost. Compared to residual GCs present in draining LNs after prime that were not boosted, we found that the frequency of GC B cells increased significantly after an ipsilateral boost due to an influx of *de novo* activated KI B cells, which could be explained by the open structure of ongoing GCs ^62,66,67^. The total number of fate-mapped KI GC B cells also increased 3- and 4-fold with low- and high affinity boosting antigen, respectively, indicating that a second immunization indeed induced further diversification of B cells that were originally activated upon prime. Interestingly however, compared to ongoing GCs that were not refueled, the fate-mapped cells of refueled GCs expressed significantly lower levels of SHMs and an increased proportion of them had no SHM. This could be due to re-entered memory B cells with lower levels of SHMs that were recruited either from circulating- or a LN-resident pool of memory B cells, as has been suggested previously ^40^. The decrease in SHM after boost was more evident with the high affinity antigen boost, in agreement with our data from the contralateral side where more memory-derived GC B cells with lower levels of SHMs were detected in secondary GCs after high affinity boost. The fact that the BCRs with the highest levels of SHM detected in fate-mapped GC B cells of residual GCs after prime were absent after high affinity boost, suggest that a high affinity boost stimulates GC B cells that bind with high affinity to leave GCs, or alternatively, to go into apoptosis ^63,64,68^. However, a more detailed investigation is needed of the refueling process, on a single GC level, since our data can be the combined result of both new secondary GCs composed of memory B cells that entered and separate, ongoing GCs from the prime, which could be regulated differently.

In conclusion, our data show that the recall response can be manipulated using different affinity antigens, where a low affinity boosting antigen rather than a high affinity antigen, promotes further diversification of memory B cells with higher levels of SHM. These results have direct implications, and could be exploited to design immunogens, for a sequential vaccination strategy against for example HIV-1, where the goal is to increase the level of SHM of specific precursor B cells with selected B cell receptors.

## Acknowledgements

We thank Tomohiro Kurosaki for the S1pr2-ERT2cre mice and Michel Nussenzweig for the 3BNC60^SI^ KI mice, the personnel at the KMA animal facility at Karolinska Institutet for assistance with the mice, the biomedicum flow cytometry core for FACS, Daniel Sheward for input on ELISA analysis, Iris Rocamonde Lago for cloning the eOD-GT8-expression vector, and Andrew McGuire for scientific discussions. We thank the National Research Council Canada for the use of pTT3-protein expression vectors. This work was funded by the ERC starting grant; VIVA 850424, a Wallenberg Academy Fellow grant, a start-up grant from the Swedish Research Council and additional funding by Karolinska Institutet.

## Supplementary figures

**Supplementary Figure 1:**
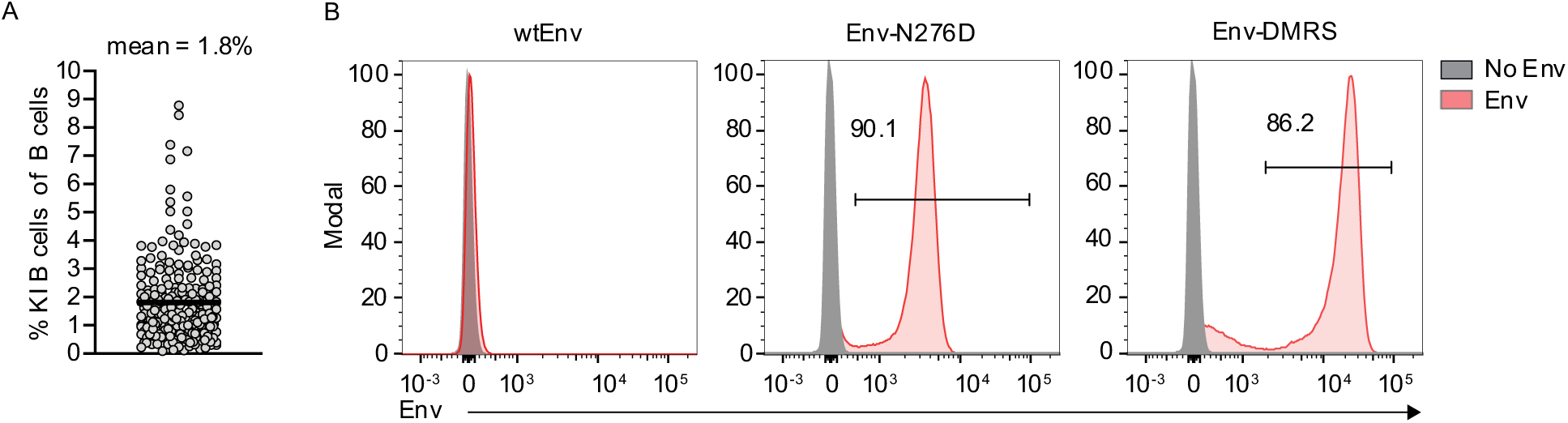
BCR expression level and binding phenotype of KI B cells used in MBMC mice. (A) Graph shows frequency KI B cells of total B cells in the blood of mixed bone marrow chimera mice 8 weeks after reconstitution. Each dot represents one mouse and data is pooled from 11 experiments. Bar represents mean value. (B) Histograms shows binding towards wtEnv, Env-N276D and Env-DMRS by Env-unstained (gray) and Env-stained (red) heterozygous KI B cells by flow cytometry.

**Supplementary Figure 2:**
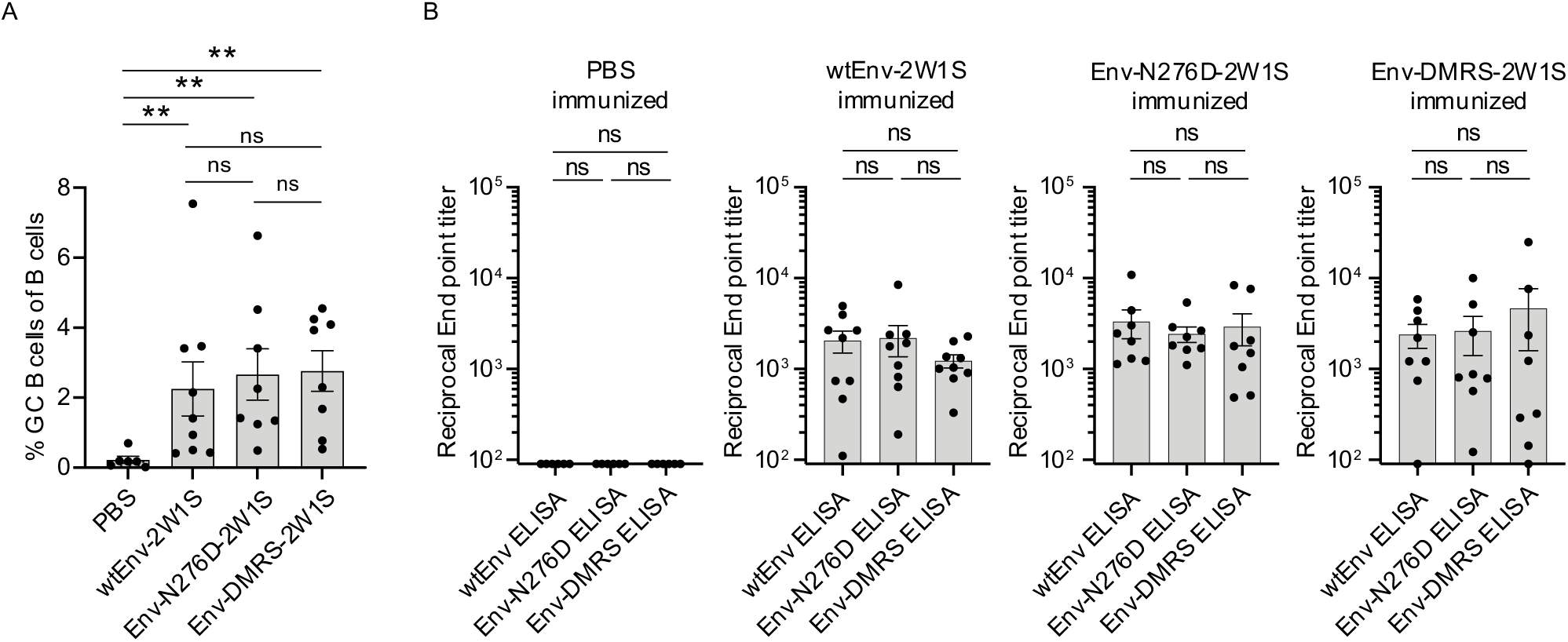
Polyclonal immune responses towards different Env antigens. Graphs show frequency of GC B cells among total B cells (A) and serum IgG responses against different Env antigens (B) of 2W1S-peptide primed C57BL/6 mice on day 15 after foot pad immunization with PBS, wtEnv-2W1S, Env-N276D-2W1S or Env-DMRS-2W1S. Each dot represents one mouse. Data is pooled from two independent experiments (n=8-9). Bars indicate me an and error bars show SEM. Statistical significance was tested using Mann-Whitney U test, unpaired t-test (A) and Wilcoxon test (B) (**, P ≤ 0.01 ; ns = non-significant).

**Supplementary Figure 3:**
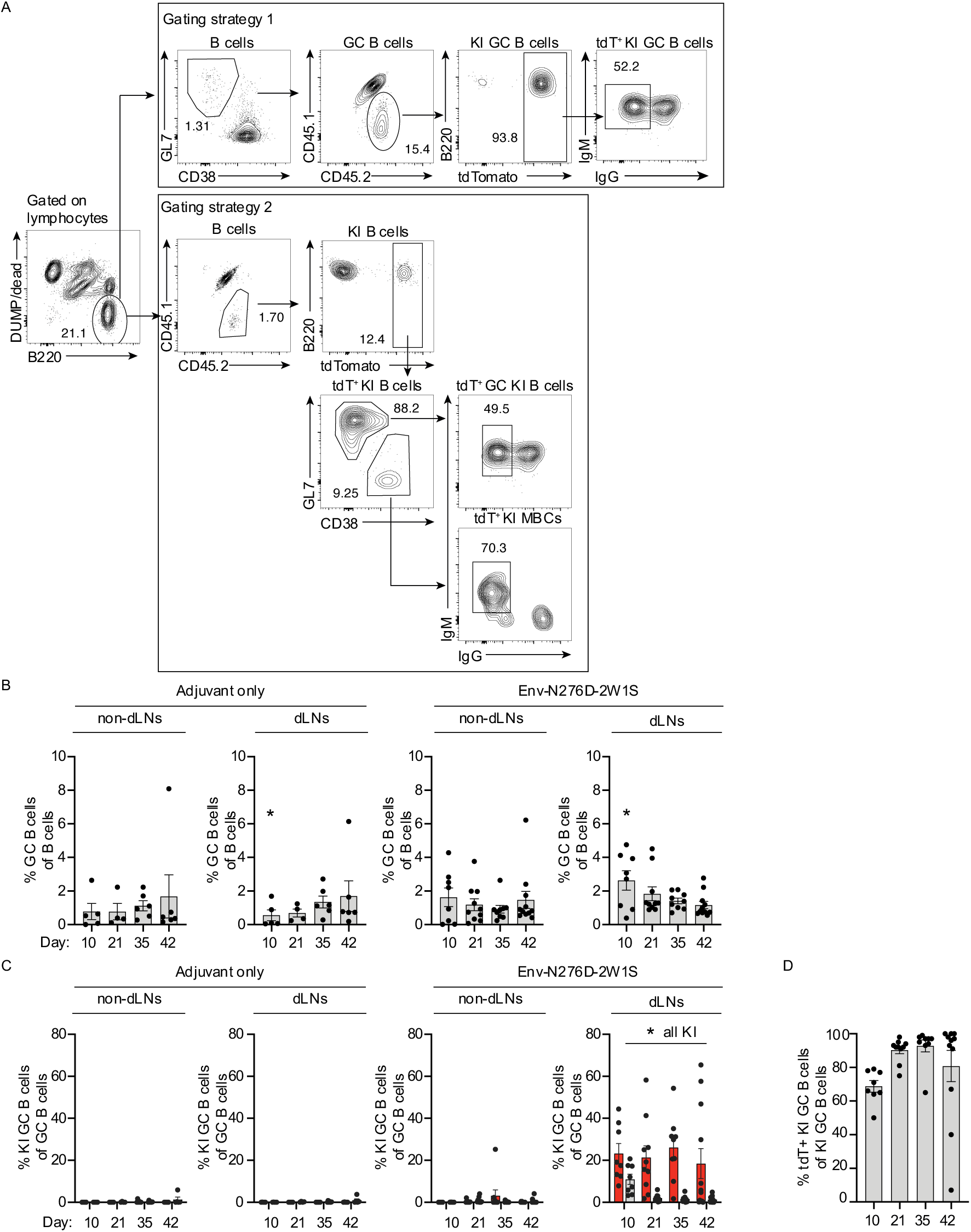
Immunization with intermediate affinity antigen and subsequent tamoxifen-treatment induces KI GC B cells of high labelling efficiency. Analysis of mixed bone marrow chimera mice treated as outlined in Figure 1A. (A) Gating strategy for analysis of the GC compartment (Gating strategy 1): total GC B cells, KI GC B cells, tdT+ KI GC B cells and for analysis of the KI memory B cell compartment (Gating strategy 2): KI B cells, activated KI B cells, KI memory B cells. (B) As in Figure 1 B but showing frequency GC B cells of B cells. (C) As in Figure 1 C but showing frequency KI GC B cells of GC B cells. (D) Graph shows tdTomato labelling efficiency of KI GC B cells over time in ipsilateral lymph nodes of mixed bone marrow chimera mice. (B-D) Each dot represents one mouse and data is pooled from at least 2 independent experiments (n= 4-29 per group). Bars indicate mean and error bars show SEM. Statistical significance was determined using Mann-Whitney U test (*, P ≤ 0.05)

**Supplementary Figure 4:**
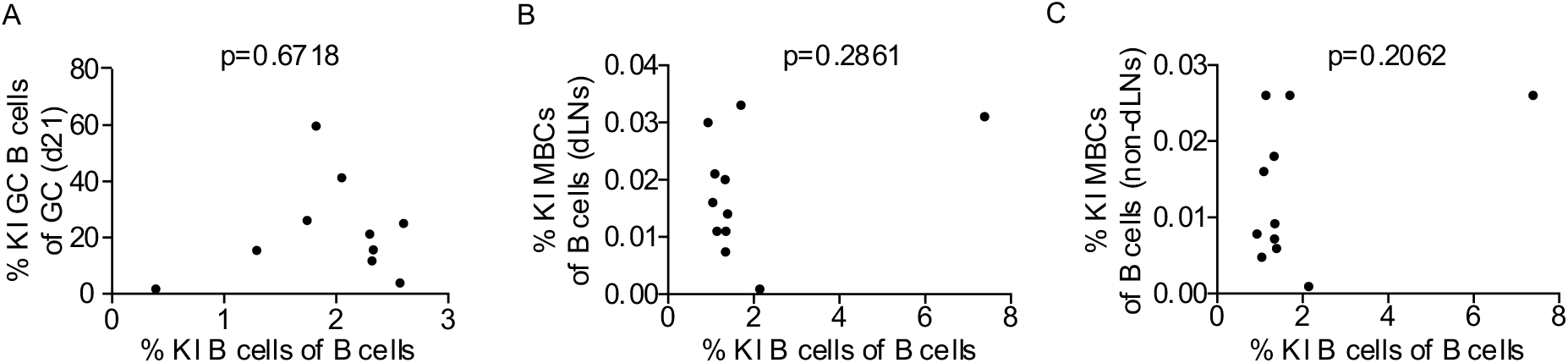
KI B cell immune response in mixed bone marrow chimera mice is not affected by KI B cell frequency. Graphs show the correlation between frequency KI B cells of B cells in blood prior to immunization and (A) frequency KI GC B cells of GC B cells at day 21 and (B) frequency KI memory B cells (MBCs) of B cells in draining lymph nodes (dLNs) and (C) non-draining lymph-nodes (non-dLNs) at day 42 post Env-N276D-immunization of mixed bone marrow chimera mice treated as in Figure 1A.

**Supplementary Figure 5:**
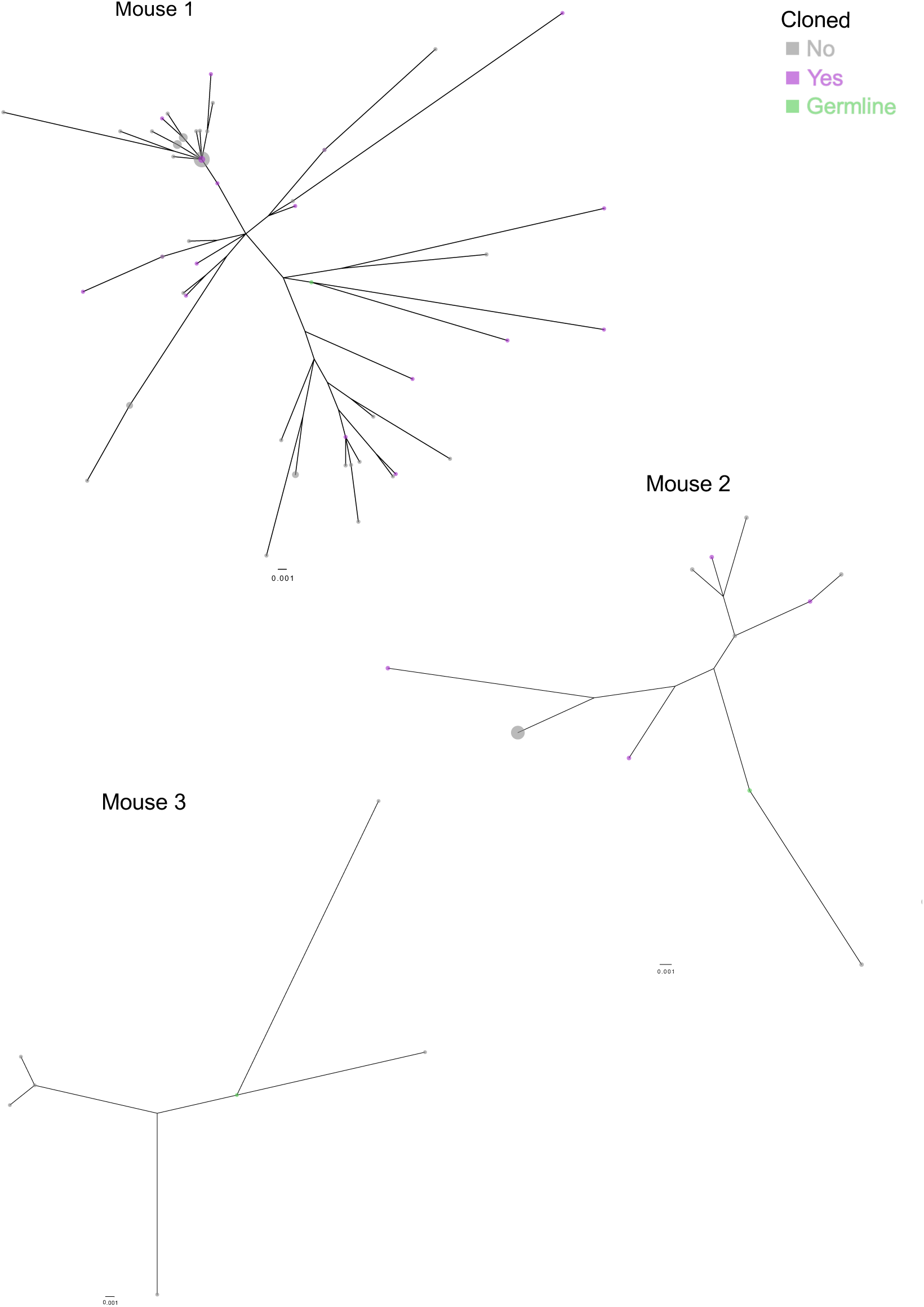
Phylogenetic trees showing the relationship of LC sequences of d42 fate-mapped KI GC B cells. Phylogenetic trees showing the relationship between light chain somatic nucleotide hypermutations of fate-mapped KI GC B sorted from draining lymph nodes 42 days after Env-N276D-2W1S immunization of MBMC mice, as shown in Figure 2C. Each tree represents 1 mouse. Trees are rooted by the germline 3BNC60 LC sequence (green). Purple dots indicate cloned BCRs shown in Figure 2E-H. Gray dots indicate non-cloned BCRs.

**Supplementary Figure 6:**
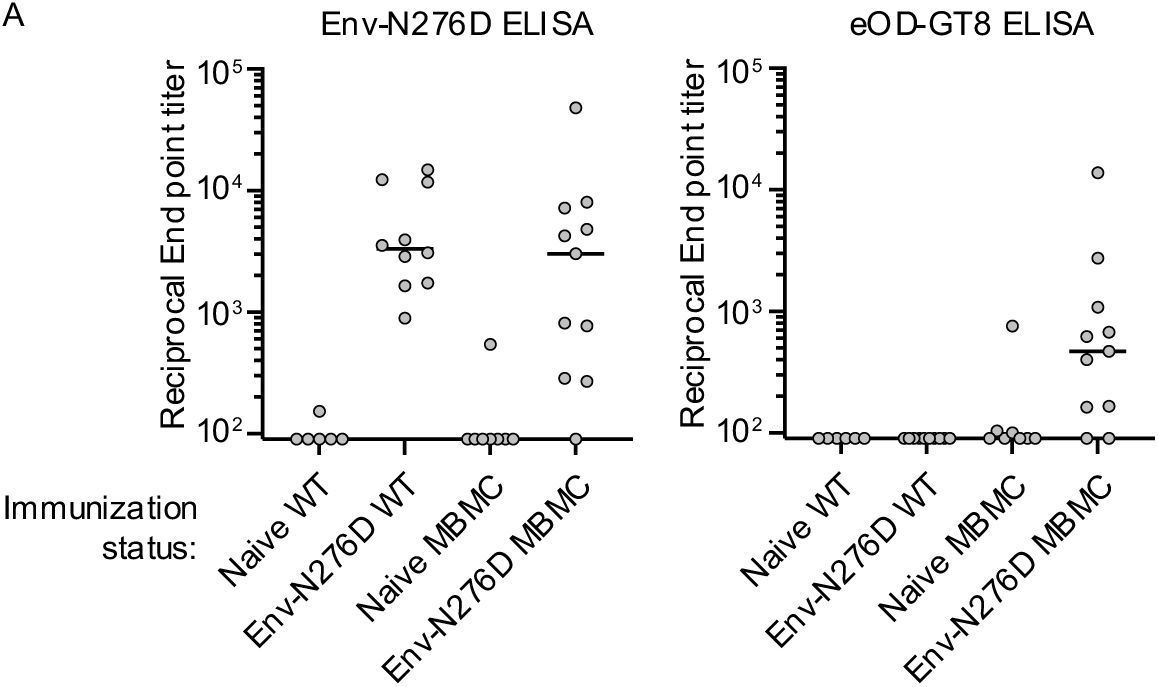
Detection of KI B cell serum antibodies by eODG-T8 ELISA. Graphs show serum antibody responses against Env-N276D (left panel) and eOD-GT8.1 (right panel) of naïve wild type (WT) mice, naïve mixed bone marrow chimera (MBMC) mice and WT and MBMC mice 15 days post footpad- and intraperitoneal immunization with Env-N276D in adjuvant. Each dot represents one mouse and data is pooled from two independent experiments (n= 6-11). Bars represent median.

**Supplementary Figure 7:**
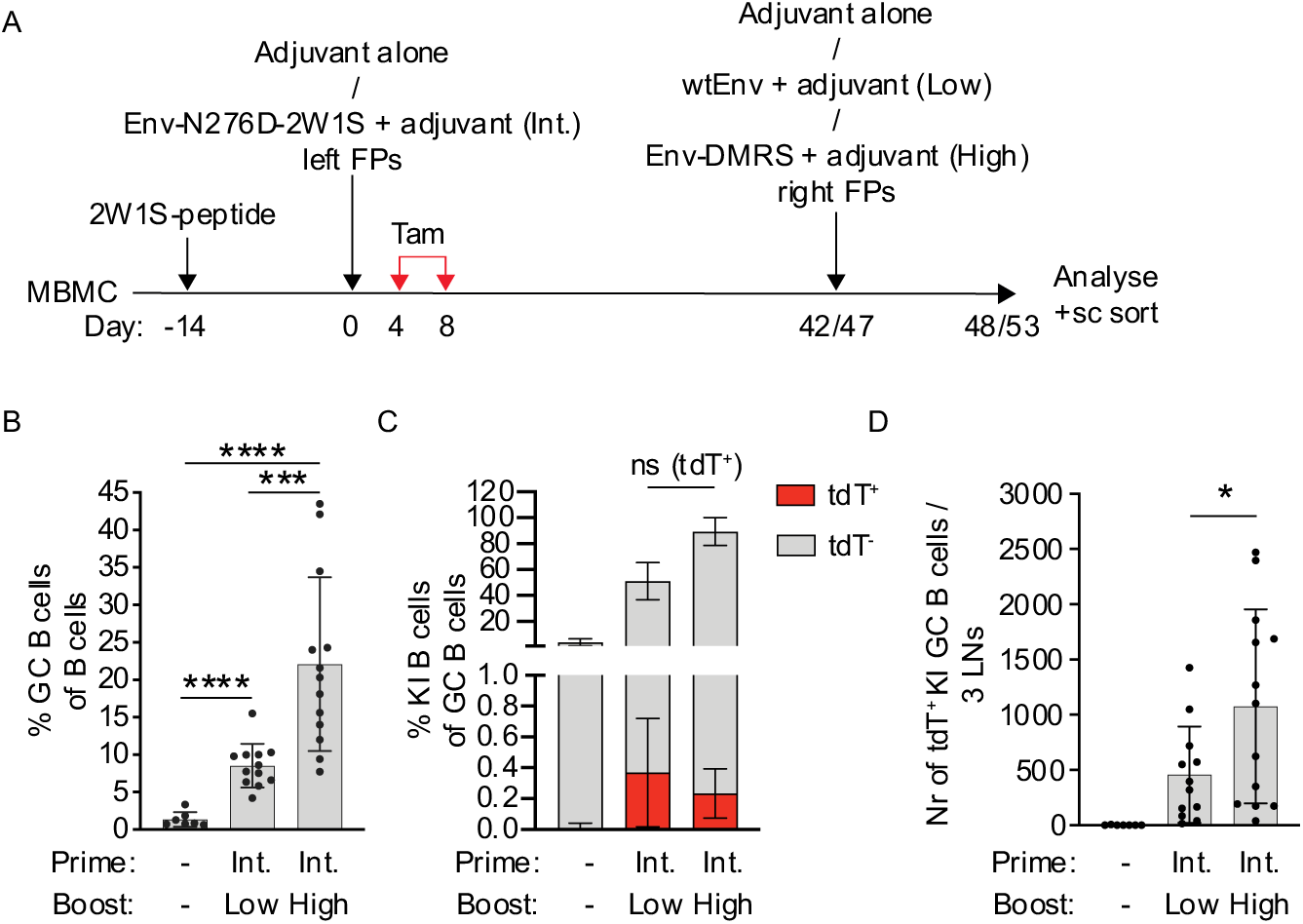
High affinity boosting antigen results in increased numbers of memory-derived GC B cells. (A) Schematic overview of the experiment: mixed bone marrow chimera (MBMC) mice were pre-primed intraperitoneally with 2W1S-peptide and immunized 2-4 weeks later in the left footpads with adjuvant only (control-immunized mice) or intermediate affinity Env-N276D-2W1S (Int.) in adjuvant. Mice were subsequently treated with tamoxifen at indicated timepoints (red arrows) and boosted 42 or 47 days post prime in the right footpads with either adjuvant only, low affinity wtEnv (Low) or high affinity Env-DMRS (High) in adjuvant. Analysis was performed 6 days post-boost. (B-D) Graphs show frequencies GC B cells (G^+^,LC7D38^-^) of B cells (B), frequencies fate-mapped (td^+^T) and non-fate-mapped (t^-^d)KTI B cells (CD45.2^+^) of GC B cells (C) and number of fate-mapped (t^+^d) TK I GC B cells per 3 contralateral lymph nodes (LNs) (D) at 6 days post boost. (C) Bars represent stacked t^-^ d(gTray) and tdT^+^ (red) KI B cell frequencies of GC B cells. Each dot represents one mouse and data is pooled from two independent experiments (n=6-13). ‘-‘ indicates immunization with adjuvant only. Bars indicate me an and error bars show SD. Statistical significance was tested using Mann-Whitney U test or Unpaired t test (*, P ≤ 0.05; ***, P ≤ 0.001; ****, P ≤ 0.0001).

**Supplementary Figure 8:**
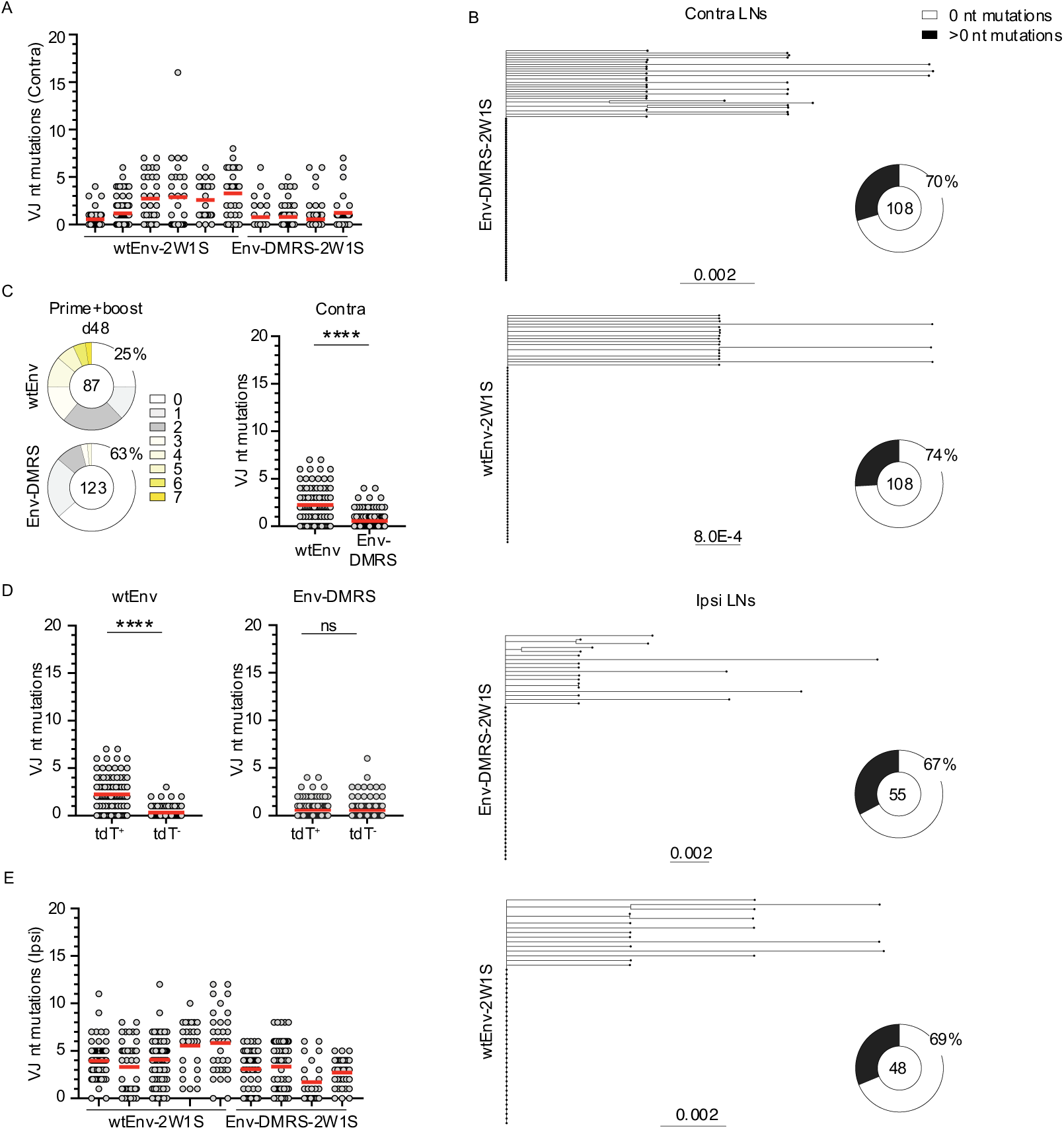
Low affinity boost results in the expansion of recalled B cells in GCs with higher levels of somatic hypermutations (A) As in Figure 5A but depicted as individual mice: graph shows number of light chain (VJ) nucleotide (nt) mutations in-antibodies of fate-mapped KI GC B cells (n= 29-85 per mouse) single cell sorted from contralateral lymph nodes 6 days after boost with wtEnv-2W1S or Env-DMRS-2W1S. (B) Pie charts show frequency of nucleotide mutations in light chain (VJ) sequences and phylogenetic trees show relationship between light chain (VJ) sequences of non-fate-mapped KI GC B cells sorted 6 days post wtEnv-2W1S or Env-DMRS-2W1S boost. Number in the pie chart indicates number of sequences analyzed. Slice size and colour indicate fraction of light chain sequences with nucleotide mutations. Percentage shows frequency of unmutated sequences. (C) Pie charts depict distribution of light chain (VJ) nucleotide mutations of fate-mapped KI GC B cells sorted from contralateral LNs 6 days after wtEnv or Env-DMRS boost of mice treated as in Supplementary figure 7A. Number in the pie chart indicates number of sequences analyzed. Slice size and colours indicate fraction of sequences with the same number of nucleotide mutations. Percentage shows frequency of unmutated sequences. Graph shows number of light chain (VJ) nucleotide mutations in antibodies of fate-mapped KI GC B cells single cell sorted from contralateral LNs 6 days after wtEnv or Env-DMRS boost of mice treated as in Supplementary Figure 7A. Data shows 1 experiment with single cells sorted from 2 mice per group (n = 87-123 sequences per group). (D) Graphs show number of light chain (VJ) nucleotide mutations in antibodies of fate-mapp^+^)evde(rtsduTs non-fate-mapped (td T^-^) KI GC B cells sorted from contralateral LNs 6 days after wtEnv (left panel) or Env-DMRS (right panel) boost of mice treated as in Supplementary Figure 7A. Data shows 1 experiment with 2 sorted mice per group (n = 82-123 sequences per group). (E) As in (A) but for ipsilateral lymph nodes. (A, C-E) Each dot represents one VJ sequence from one cell. Red bars indicate mean values. Statistical significance was tested using Mann-Whitney U test ****, P ≤ 0.0001).

**Supplementary Figure 9:**
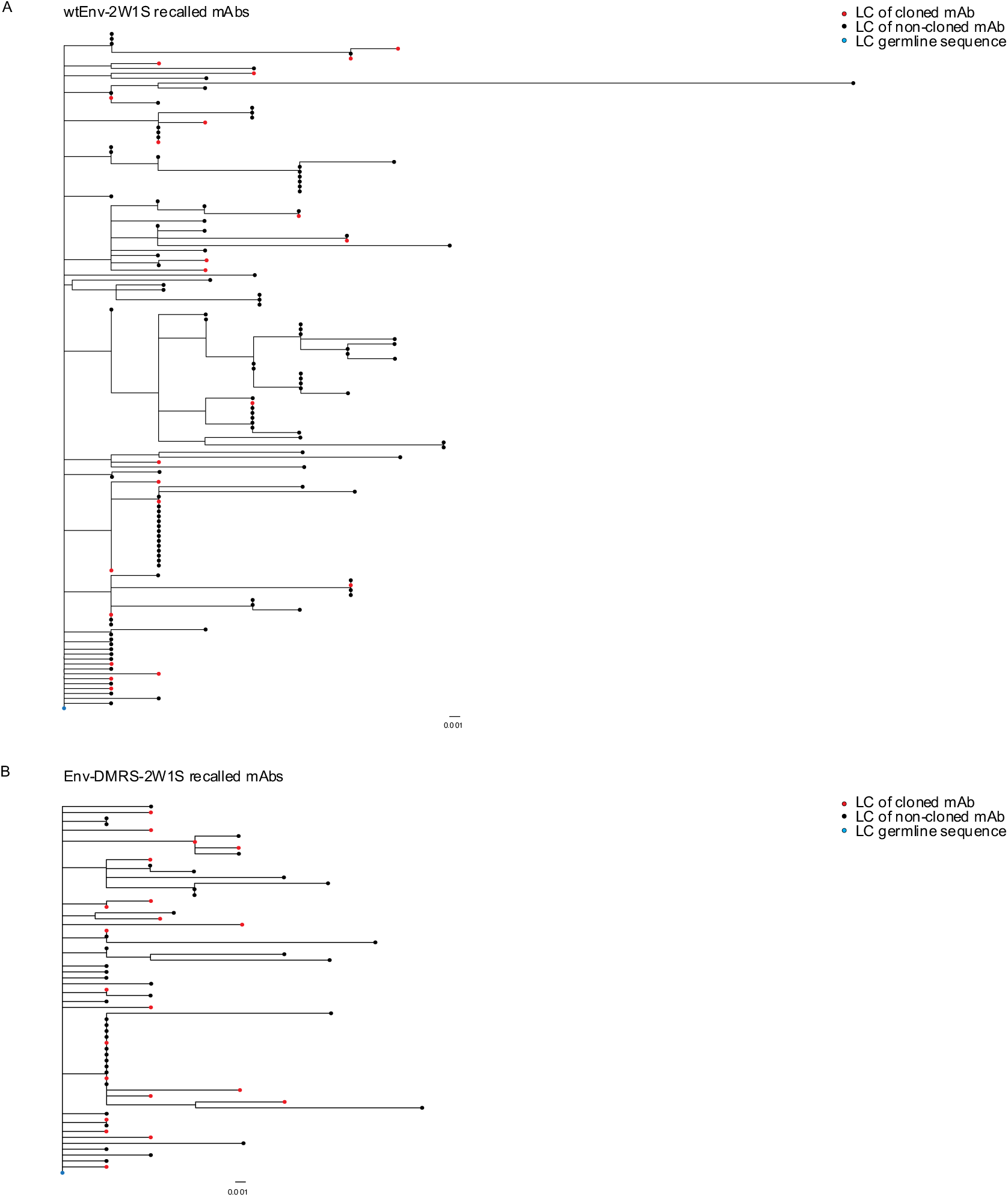
Phylogenetic trees showing the relationship of sequences of memory-derived GC B cells. Phylogenetic trees showing the relationship between VJ nucleotide mutations of fate-mapped KI GC B sorted from contralateral LNs 6 days after wtEnv-2W1S or Env-DMRS-2W1S boost of mice treated as in Figure 3A. Trees are rooted by the germline 3BNC60 VJ sequence (blue dot). Red dots indicate cloned BCRs shown in Figure 5C-E.

